# The *Bcvic1* and *Bcvic2* vegetative incompatibility genes in *Botrytis cinerea* encode proteins with domain architectures involved in allorecognition in other filamentous fungi

**DOI:** 10.1101/2022.04.03.486892

**Authors:** Saadiah Arshed, Murray P. Cox, Ross E. Beever, Stephanie L. Parkes, Michael N. Pearson, Joanna K. Bowen, Matthew D. Templeton

**Affiliations:** Bioprotection, New Zealand Institute of Plant and Food Research, Auckland, New Zealand; School of Biological Sciences, University of Auckland, Auckland New Zealand; Bioprotection Aotearoa Centre of Research Excellence, New Zealand; School of Natural Sciences, Massey University, Palmerston North, New Zealand; Maanaki Whenua Landcare Research, Auckland, New Zealand

## Abstract

Vegetative incompatibility is a fungal allorecognition system characterised by the inability of genetically distinct conspecific fungal strains to form a viable heterokaryon, and is controlled by multiple polymorphic loci termed *vic* (*v*egetative *i*n*c*ompatibility) or *het* (*het*erokaryon incompatibility). We have genetically identified and characterised the first *vic* locus in the economically important, plant-pathogenic, necrotrophic fungus *Botrytis cinerea*. A bulked segregant approach coupled with whole genome Illumina sequencing in near-isogenic lines of *cinerea* was used to map a 60-kb genomic region for a *vic* locus. Within that locus, we identified two adjacent, highly polymorphic open reading frames, *Bcvic*1 and *Bcvic*2, which encode predicted proteins that contain domain architectures implicated in vegetative incompatibility in other filamentous fungi. *Bcvic*1 encodes a predicted protein containing a putative serine esterase domain, a NACHT family of NTPases domain, and several Ankyrin repeats. *Bcvic*2 encodes a putative syntaxin protein containing a SNARE domain; such proteins typically function in vesicular transport. Deletion of *Bcvic*1 and *Bcvic*2 individually had no effect on vegetative incompatibility. However, deletion of the region containing both *Bcvic*1 and *Bcvic*2 resulted in mutant lines that were severely restricted in growth and showed loss of vegetative incompatibility. Complementation of these mutants by ectopic expression restored the growth and vegetative incompatibility phenotype, indicating that *Bcvic*1 and *Bcvic*2 are controlling vegetative incompatibility at this *vic* locus.

**Author Summary:** Fungal colonies are characterised by radiating filaments, termed hyphae, which often fuse to form a highly interconnected individual. This is advantageous since it enables efficient water and nutrient utilisation across a colony network. However, hyphal fusion is not necessarily restricted to within an individual colony, with potential for hyphal fusion between individuals belonging to the same species. There are, however, drawbacks to this. For instance, viruses that detrimentally affect a colony may be transmitted, with their infection leading to a reduction in the virulence of a pathogenic species. Fungi have therefore developed complex systems to prevent fusion between genetically distinct individuals of the same species. This phenomenon is termed vegetative incompatibility and results in the death of fused cells and cessation of transfer of cellular contents from one individual to another. We have identified the first genes in the fungal plant pathogen *Botrytis cinerea* that control this phenomenon. They resemble genes that control vegetative incompatibility in other fungi, and genes involved in immunity in plants and animals. Uncovering further genes involved in vegetative incompatibility in *B. cinerea* may pave the way for the development of a ‘super donor’ strain capable of overriding vegetative incompatibility to transmit viruses, thus enabling their exploitation as potent control agents against this damaging plant pathogen.

## Introduction

Cell fusion is common to all eukaryotic kingdoms of life. It is perhaps most readily apparent in fungi, where a mycelium is formed by the fusion (anastomosis) of hyphal filaments during vegetative (asexual) growth, giving rise to a syncytium-like complex of interconnected cells operating as a coordinated individual. This allows the flow of organelles and cytoplasm between hyphal compartments, serving a number of functions that are beneficial to the fungal colony. These include the translocation of water and transport of nutrients, thereby regulating overall homeostasis across a wide range of nutrient limiting spatial scales [1], and enabling efficient utilisation of a nutrient source [2, 3]. Cell fusion in fungi can also drive the transition from unicellularity to multicellularity [4], with fusion between identical genotypes improving fitness of the developing colony [2, 3].

However, there are downsides to fungal cell fusion, especially between genetically non-identical individuals. Cytoplasmic mixing can enable the transmission of deleterious mitochondrial genotypes, which depress fitness [5, 6], and pathogens (e.g. mycoviruses [7–9]). Allorecognition (the ability to discriminate between self and conspecific non-self) is therefore fundamental for survival, since it can preclude cell fusion between non-identical individuals and potentially deleterious fitness consequences. Most genetically distinct fungal individuals from the same species are unable to anastomose. Indeed, the model fungus *Neurospora crassa* has evolved multiple mechanisms to avoid cell fusion between genetically distinct individuals at all costs [10].

The multifaceted *N. crassa* allorecognition system is far from unique; it is becoming increasingly apparent that mechanisms evolved in fungi to avoid cell fusion are complex, often comprising multiple checkpoints. These checkpoints, at which successful fusion can be arrested, operate either pre-fusion (curtailing chemotropism [11] and preventing cell wall fusion [12]) or post-fusion (involving intracellular recognition in germlings and/or mature hyphae [13–16]). Post-fusion mechanisms following allorecognition involve the triggering of programmed cell death (PCD) of the heterokaryotic fusion cell, and subsequent restoration of the two separate fungal entities, a process termed vegetative (VI) or heterokaryon incompatibility (HI) [16–18].

VI is initiated in hyphae when genetically distinct individuals from the same species differ at specific loci termed *v*egetative *i*n*c*ompatibility (*vic*) or *het*erokaryon incompatibility (*het*) loci [17, 19–21]. Genetic and modelling analyses indicate that filamentous fungi typically have 6–12 *vic* loci [22, 23], but may have as many as 30 [16, 24, 25].

Incompatibility systems have been described in numerous divisions of filamentous fungi including ascomycete, basidiomycete and glomeromycete species [21, 26, 27]. However, the molecular mechanisms of the VI phenomenon have largely been identified in a small number of well-studied systems, notably *N. crassa*, *Podospora anserina* and *Cryphonectria parasitica*, which all belong to the Sordariomycetes (reviewed in [16]). Loci involved in VI are typically highly polymorphic and found in hypervariable genomic regions [28]. VI can involve allelic genes, for example, the *het-c* incompatibility loci in *N. crassa* [29], or be controlled by a non-allelic system, for example, the *het-c/het-d/het-e* incompatibility loci in *P. anserina* [30–32]. The polypeptides encoded by the three *het-c* alleles in *N. crassa* are similar except for a variable domain of 34–48 amino acids. This polymorphic region is sufficient to confer *het-c* allelic specificity. *het-c* is tightly linked with the *pin-c* (*p*artner for *in*compatibility) locus which encodes a protein with a conserved domain termed HET [29]. In contrast with *het-c*, the three *pin-c* alleles are highly polymorphic (∼55% amino acid identity) in the region outside the conserved HET domain. Genetic interactions between different *het-c* and *pin-c* alleles (e.g. *het-c1* and *pinc-2*) result in allorecognition, with allelic interaction between different *het-c* alleles (e.g. *het-c1* and *het-c2*) increasing the severity of the incompatible reaction. The *het-c* locus of *P. anserina* encodes a glycolipid transfer protein [31], which interacts with the encoded proteins of *het-d* and *het-e*. *het-d* and *het-e* are paralogues which belong to a large gene family. The encoded proteins are tri-partite nucleotide oligomerisation domain (NOD)-like receptors (NLRs) [30, 32], which in the case of *het-d* and *het-e* comprise an N-terminal HET domain, a central NACHT domain with a nucleotide-binding site and a C-terminal WD40 repeat domain.

In addition to *vic* genes, other loci are involved in the initiation of a programmed cell death (PCD) response following allorecognition post cell fusion, which are not classified as *vic* or *het* loci since they can regulate germling-regulated death (GRD) in addition to hyphal PCD. These have been identified in *N. crassa*: *sec-9*/*plp-1* [13] and *rcd-1* [14], but are also present in other fungi [13]. The *sec-9*/*plp-1* system relies on the physical interaction of the PLP-1 patatin-like phospholipase-1 NLR protein, which comprises an N-terminal patatin-like phospholipase domain, a central nucleotide-binding domain and a C-terminal tetratricopeptide repeat (TPR), with a SEC-9 protein, which is characterised by a soluble N-ethylmaleimide-sensitive factor attachment protein receptor (SNARE) domain of different specificity [13]. In contrast, *rcd-1* belongs to a highly polymorphic gene family with coexpression of antagonistic alleles resulting in PCD [14].

The allorecognition system in fungi can therefore be viewed as modular, with a recognition module comprised of highly polymorphic protein domains that act as the specificity region required for recognition, and a death effector module that induces the compartmentalisation and cell death process [33]. These modules can either be present within a single protein or on different protein components of the allorecognition system [16]. The repeat domains of NLR-like proteins, if involved, are postulated to play a role in recognition of the specificity determinants, whilst the HET domain, which is specific to filamentous fungi and present in most proteins encoded by *vic* genes, is thought to be involved in the initiation of the cell death reaction.

How the interactions between alternate incompatibility proteins translate into cell death is not well understood. Identification of the genes involved has, generally, suggested mechanisms for cell death induction [16]. For example, the phospholipase domain of PLP-1 may directly alter membrane phospholipids, with the PLP-1 complex acting as a membrane toxin itself [13]. However, clear evidence of mechanism has been demonstrated only for the *het-s*/*het-S* system in *P. anserina* [34, 35]. *het-s* encodes a prion protein. Depending on the epigenetic state of the strain, the protein is either monomeric and soluble [Het-s*], or assembles into prion aggregates [Het-s]. Het-s consists of two domains: an N-terminal globular domain (HeLo) and a C-terminal prion-forming domain (PFD). Het-S has a similar domain structure but cannot form a prion. Interaction between the prion form of Het-s and Het-S results in a conformational change in the HeLo domain of Het-S, leading to acquisition of pore-forming activity and relocation to the cell membrane where it compromises membrane integrity [36–39].

In *P. anserina*, the cell death reaction is characterised downstream by the induction of a specific set of genes (*idi*), followed by the production of numerous degradative enzymes including proteases, laccases and phenoloxidases, increased vacuolisation, increased deposition of septa, accumulation of lipid droplets and the abnormal deposition of cell wall material [33, 40, 41]. Along with these hallmarks of cell death, autophagosomes are observed in the cytoplasm soon after the triggering of cell death by incompatibility. Electron microscopy has confirmed the double plasma membrane present in the autophagosome, and its cytoplasmic content, which are characteristic of a type II programmed cell death pathway. It is hypothesised that the induction of autophagy may function to control PCD by protecting neighbouring cells from cell death [42]. In *C. parasitica*, VI is associated with the accumulation of secondary metabolites and reactive oxygen species (ROS), with pheromone processing hypothesised to be an important component of allorecognition since crypheromonins, which resemble type ‘a’ mating pheromones, accumulate [43, 44].

Whilst having implications for the survival of fungal colonies, VI is also of relevance when formulating biological control strategies involving mycoviruses [45]. The use of hypovirulent mycoviruses as biological control agents (BCAs) requires that they either be mechanically transmissible, which would necessitate multiple applications to control disease, or able to infect the entire population of a fungal phytopathogen, by being readily transmissible from one isolate to another. However, VI can limit the efficacy of the latter class of potential hypovirulence-inducing mycoviruses. For example, transmission of the hypovirulent mycovirus that infects *C. parasitica*, the casual agent of Chestnut blight, has been limited by VI, in turn limiting the use of the virus as a BCA [7].

*Botrytis cinerea* Pers. Fr. (teleomorph *Botryotinia fuckeliana* (de Bary) Whetzel) is an economically important plant-pathogenic fungus that causes grey mould disease in over 280 mainly dicotyledonous species [46, 47]. *B. cinerea* is a necrotrophic pathogen that first colonises necrotic, senescent or dead tissue, largely as a saprobe. From this base, it is capable of infecting live tissue, often through the production of an infection cushion [48]. Mycoviruses which result in a hypovirulent phenotype have been identified in *B. cinerea* [e.g. 49, 50, 51]. These could be effective as BCAs although the VI system in *B. cinerea*, as with other fungal systems, may preclude their efficacy. Super donor strains or combinations of strains have been developed in *C. parasitica*, using gene disruption of *vic* loci combined with classical genetics, enabling transmission of a hypovirulent mycovirus independent of *vic* genotype [52, 53]. Thus in addition to being of interest with regard to the ecology of a damaging fungal pathogen and the evolution of fungal fitness, identifying the molecular components of VI in *B. cinerea* may aid the development of effective BCAs for control of this damaging pathogen. Beever and Weeds [54] confirmed the existence of 66 *v*egetative *c*ompatibility *g*roups (vcg) from single ascospores within field isolates of *B. cinerea*. Population genetic analysis of those vcgs indicated the presence of at least seven *vic* loci in New Zealand *B. cinerea* isolates [54]. This paper describes the identification of the first *vic* locus in *B. cinerea*.

## Results

### The F1BC3 near-isogenic lines segregated 1:1 at the BenA and *vic* loci

The F1BC3 progeny of *B. cinerea* used in this study were generated from three generations of backcrosses to the recurrent parent REB749-8 (S1 Fig.). The backcrossing strategy was deemed to be complete when there was 1:1 segregation of the benomyl (BenA) resistance and *vic* phenotypes. Fifteen out of 32 progeny were found to be sensitive to benomyl and the other 17 resistant (47% B^R^; 53% B^S^). As expected, all the progeny in the F1BC3 generation displayed ultra-low level dicarboximide resistance when plated onto Malt Extract Agar (MEA) + carbendazim (100 mg/L) since both parents carried the same allele for ultra-low level dicarboximide resistance. Following mycelial compatibility testing using nitrate non-utilising mutants, 18 progeny were compatible with REB749-8 and incompatible with REB811-28 and bulked as vcg1, whereas 14 progeny from the REB839 series were incompatible with the recurrent parent REB749-8 and compatible with the non-recurrent F1BC2 parent REB811-28, and were bulked as vcg2. Two isolates from the REB839- population, REB839-5 and REB839-1, were used as the near-isogenic vegetative incompatibility test strains, vcg1 and vcg2, and for functional characterisation of candidate *vic* genes through transformation (S1 Table).

### A candidate *vic* locus lies on scaffold 56 of the *B. cinerea* T4 reference genome

Approximately 14 and 18 million read pairs were obtained for the genomes bulked in the vcg1 and vcg2 samples, respectively. When the reads were mapped to the 118 scaffolds of the *B. cinerea* T4 reference sequence there was an average of 43 and 62 reads per nucleotide position for the vcg1 and vcg2 bulks, respectively, and the genome coverage that was shared between both vcg1 and vcg2 was 92.4% (36,497,613 bp) of the T4 genome. Not all the genome was covered owing to the strict read-mapping thresholds used to ensure highly accurate variant calling. The missing regions consisted of repetitive elements such as microsatellites and transposons. Furthermore, a small proportion of reads that had a high divergence in sequence identity with respect to the reference did not align and therefore were excluded from the analysis.

The single nucleotide polymorphism (SNP) profiles identified from the vcg1 and vcg2 bulks fell into three groups. The most abundant type were those SNPs that were shared between both bulks that were identical in all the progeny within each bulk, reflecting the expected considerable regions of isogeny within the backcrossed offspring with no linkage to the trait of interest. The second category of SNPs were those shared between bulks with some variability within each individual bulk, indicative of the non-isogenic genomic regions in the parents, with random segregation of these SNPs in the progeny, with no linkage to the VI phenotype used to differentiate the bulks. The vcg1 and vcg2 bulks shared 139,759 SNPs that were uniformly distributed throughout the genome relative to the T4 reference.

The third and most important class of SNPs were the bulk-specific SNPs that were present in one bulk but absent in the other. Initially 759 vcg1 and 1,478 vcg2 bulk-specific SNPs were identified using the SNP-calling algorithm, revealed by applying a 5000-bp sliding window with a 25-bp lag to the entire genome sequence. These bulk-specific SNPs were tightly clustered within a few scaffolds (Fig. 1). The vast majority of these were discarded from further analysis following manual curation. As expected, most of the SNPs occurred within microsatellites or homopolymer runs, which can cause problems during read mapping. A proportion were miscalled insertions and deletions, whilst others were simple sequencing errors. Miscalled bulk-specific SNPs often occurred because of the strict SNP-calling thresholds that were applied, whereby SNPs needed to be represented by at least eight reads in one bulk and absent in the other. For instance, a SNP that was shared or heterogeneous between vcg1 and vcg2 could be miscalled as a bulk-specific SNP by the SNP-calling algorithm if the alignment for either bulk had a region of low sequence coverage represented by less than eight reads. After the above analysis had removed the probable spurious differential SNP calls, the only T4 reference genome scaffold that had large numbers of accurate bulk-specific SNPs was bt4_SupSuperContig_110r_56_1 (scaffold 56; 449,055 bp long; this corresponds to the B05.10 region 143,339 to 203,859 on scaffold 1.28). There were 307 and 369 bulk-specific SNPs in the vcg1 and vcg2 bulks across this scaffold, with over 98% of them in close proximity (300 of the 307 with respect to the vcg1 bulk and all of those identified with respect to the vcg2 bulk), with each SNP within at least 5000 bases of another. In certain regions of scaffold 56, there were occurrences of more than 60 bulk-specific SNPs within a 5-kb window (S2 Fig.). A region approximately 60 kb in length on scaffold 56, from position 143,443 to 204,125, had the highest density of bulk-specific SNPs, and was selected as a candidate *vic* gene locus for further interrogation (Fig. 2).

**Fig. 1.**
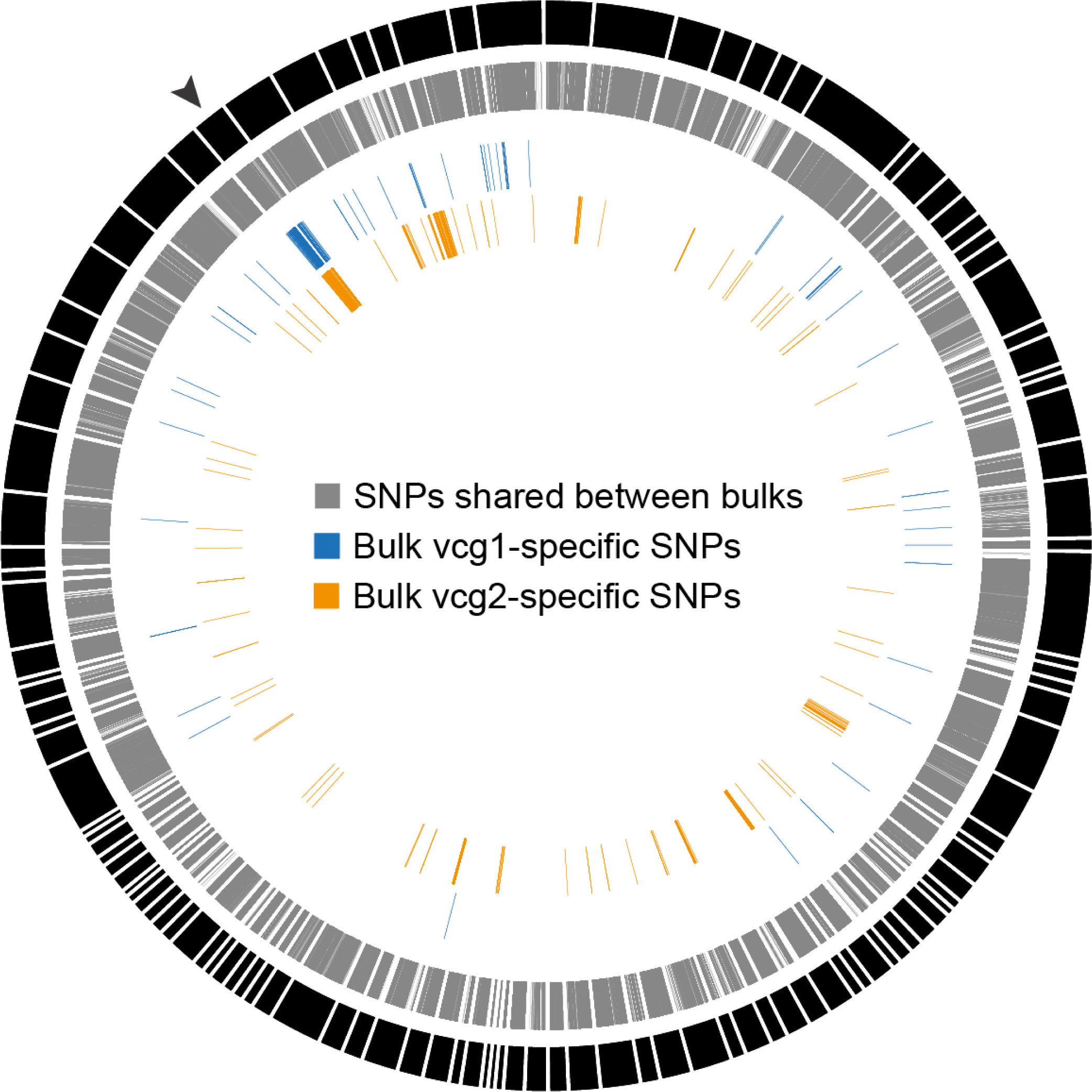
Distribution of Single Nucleotide Polymorphisms along the *Botrytis cinerea* T4 genome. The outer black bars show the 118 scaffolds that make up the *B. cinerea* T4 reference genome. The dense, uniformly distributed grey bars represent the large number of Single Nucleotide Polymorphisms (SNPs) shared between the vcg1 and vcg2 bulks. The outer blue and inner orange lines represent vcg1 and vcg2 bulk-specific SNPs, respectively, which are in close proximity to each other, occurring within 5 kb of another bulk-specific SNP. The black arrow denotes the location of scaffold bt4_SupSuperContig_110r_56_1. Note the preponderance of bulk-specific SNPs at this location.

**Fig. 2.**
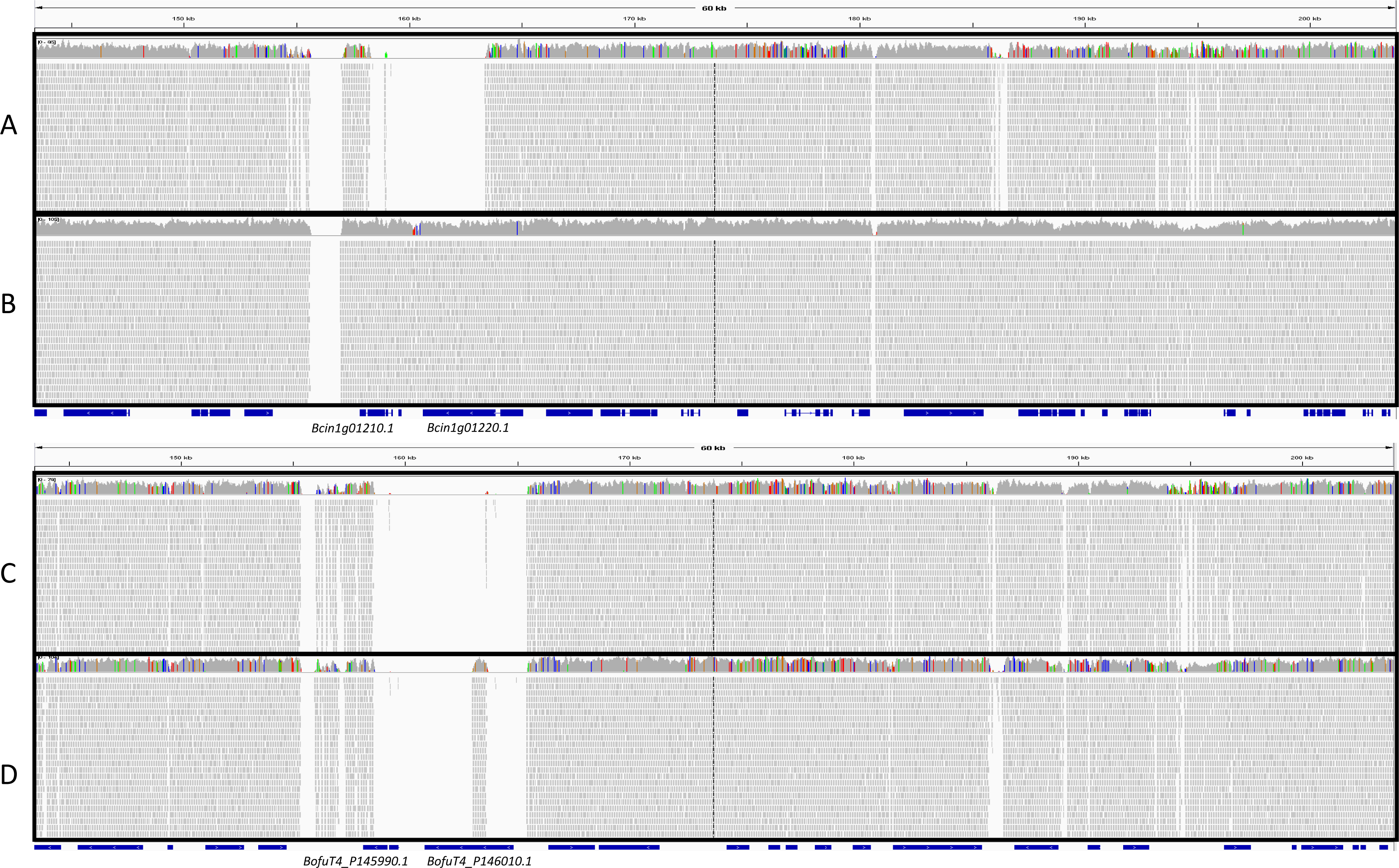
Mapping of bulked vcg1 and vcg2 reads to the T4 and B05.10 *Botrytis cinerea* reference genomes at the candidate *vic* locus. (A, B) Bulked vcg1 (A) and vcg2 (B) reads mapped to B05.10. The region shown includes the scaffold positions 143,339 to 203,859 on scaffold 1.28. (C, D) Bulked vcg1 (C) and vcg2 (D) reads mapped to the T4 reference showing the scaffold positions 143,443 to 204,125 on scaffold bt4_SupSuperContig_110r_56_1 (scaffold 56). Gene predictions are shown by the blue bars. Sequencing reads are denoted by the grey bars within each panel. Single nucleotide polymorphisms (SNPs) are indicated by the different colours in the coverage panel above the sequencing reads.

### Two genes at the candidate *vic* locus share similarities with previously identified fungal *vic* genes

Twenty-four genes were located in the candidate region on scaffold 56 of the T4 genome, but only fifteen had orthologous gene predictions on chromosome 1 of the gapless B05.10 genome, casting doubt on the reliability of nine of the T4 gene models. Of the remaining fifteen predicted genes, six had 100% amino acid identity between vcg1 and vcg2 bulks and were therefore excluded from further analysis. The remaining nine candidate genes are listed in S2 Table. Three of these genes had no putative predicted function since no domains were identified following analysis with Pfam and Interproscan. Of the remaining genes, Bcin1g01210.1 and Bcin1g01220.1 from the B05.10 genome (equivalent to BofuT4_P145990.1 and BofuT4_P146010.1 in the T4 genome, respectively), encoded proteins with domains consistent with a role in allorecognition. In addition, both had missing sequence alignment coverage when the sequences from both bulks were compared with the T4 genome, and when vcg1 sequences were compared with the B05.10 reference genome, suggesting a high degree of polymorphism (a known trait of *vic* genes). Of note is that the vcg2 bulk shared identical sequences with the gapless B05.10 reference genome for all the predicted genes in the region (Fig. 2B). When the sequence of the genes and predicted proteins were compared following polymerase chain reaction (PCR) amplification, they had a significantly lower sequence identity than the other candidates (S2 Table).

### Bcin1g01220.1 (BofuT4_P146010.1): Bcvic1

The first candidate gene, *Bcin1g01220.1* (*BofuT4_P146010.1*), henceforth referred to as *Bcvic1*, and thus *Bcvic1-1* in the vcg1 tester strain 839-5 (with the additional *-1* referring to the vcg), had a transcript length of 4,215 bp with four exons when confirmed by PCR amplification, cloning and re-sequencing, and a predicted translated protein of 1,404 amino acids. *Bcvic1* in vcg2 (*Bcvic1-2*) was 100% identical at the nucleotide and predicted amino acid sequence levels with the B05.10 sequence. The *Bcvic1* gene, as confirmed by PCR, cloning and Sanger sequencing, from vcg2 and B05.10 (*Bcvic1-2*) had a transcript length of 4,489 bp with three exons and the predicted translated protein was 1,444 amino acids long.

The percentage identity between *Bcvic1-1* and *Bcvic1-2* at the nucleotide level was 68.3% over the entire sequence. The 5′ end of the gene from nucleotide position 1 to 1,662 was more conserved, with a percentage pairwise identity of 99%, than the downstream region from position 1,663 to 4,589, which was variable, with a pairwise identity of only 50.7%. The two predicted protein sequences encoded by *Bcvic1-1* and *Bcvic1-2* contained a putative serine esterase/lipase domain (residues 52-179 in both BCVIC1-1 and BCIVC1-2; Pfam: PF05057, serine esterase; Interproscan: IPR007751: Domain of unknown function found within a group of putative lipases, including the phospholipase B YOR059C (Lpl1) from budding yeast), a NACHT domain (Residues 224-350 in BCVIC1-1 and 363-489 in BCIVC1- 2; Pfam: PF05729), and ankyrin repeats (residues 733-805, 813-903, 973-1055, 1103-1189, 1201-1285 in BCVIC1-1 and 866-944, 953-1042, 1114-1196, 1207-1231, 1241-1328, 1339-1426 in BCIVC1-2; Pfam: PF00023) (Fig. 3). The proteins had a percentage identity of 60.2% over the entire sequence. The N-terminal region from position 1 to 523, which contains the putative serine esterase and NACHT domains, shares a much higher percentage identity, of 98.7%, than the ankyrin repeat containing region downstream of the NACHT domain, which is more variable, with a percentage pairwise identity of 38.7%.

**Fig. 3.**
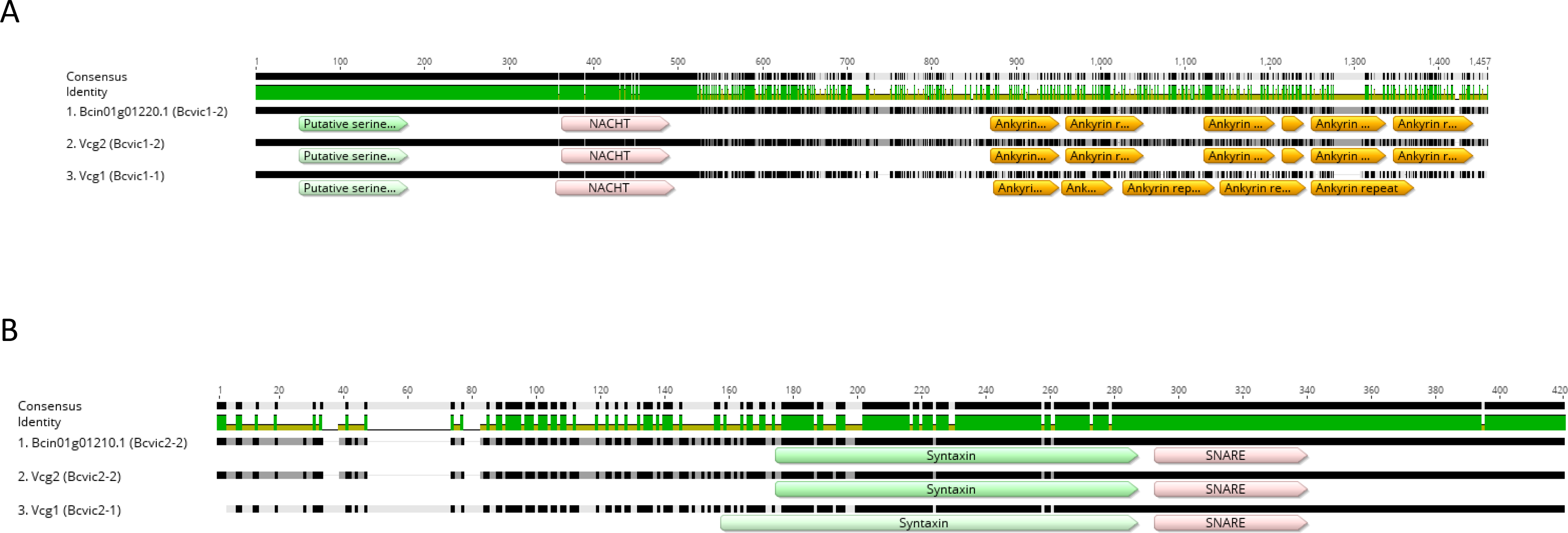
Amino acid alignments of the candidate *vic* proteins BCVIC1 (A) and BCVIC2 (B) in the *Botrytis cinerea* strains 839-5 (vcg1), 839-1 (vcg2) and B05.10. The numbers on top of the figures represent the position of the amino acid residues in the sequence alignment. The black shading in the alignment indicates that the residue at that position is the same across both sequences (100% identity); grey represents less than complete identity; and white represents very low identity for a particular position. (A) The green, pink and orange blocks denote the positions of the putative serine esterase, NACHT and ankyrin repeat domains in BCVIC1, respectively. (B) The green and pink blocks denote the positions of the syntaxin and SNARE domain in BCVIC2, respectively.

### Bcin1g01210.1 (BofuT4_P145990.1): Bcvic2

The sequence of the second candidate gene, *Bcin1g01210.1* (*BofuT4_P145990.1*) henceforth referred to as *Bcvic2* in the vcg1 and vcg2 tester strains 839-5 and 839-1, was confirmed by PCR amplification, cloning and re-sequencing. The *Bcvic2* gene from vcg1 (*Bcvic2-1*) had a transcript length of 1,254 bp, with three coding exons. The predicted translated protein was 417 amino acids long. The *Bcvic2* gene from vcg2 (*Bcvic2-2*) had a transcript length of 1,158 bp, with four coding exons. The predicted translated protein was 385 amino acids long. The sequences for *Bcvic2* in vcg2 and B05.10 were identical.

The percentage identity between *Bcvic2-1* and *Bcvic2-2* at the nucleotide level was 76.3% over the entire sequence alignment. The 5′ end of the gene from nucleotide position 1 to 931 was more variable than the 3′ end, with a percentage pairwise identity of 65.1%, contrasting with the 97.9% identity of the downstream region, between positions 932 and 1416. The two predicted protein sequences encoded by *Bcvic2-1* and *Bcvic2-2* contained a syntaxin domain (residues 155-284 in BCVIC2-1 and 140-251 in BCIVC2-2; Pfam: PF00804) and a SNARE domain (residues 290-337 in BCVIC2-1 and 257-304 in BCIVC2-2; Pfam: PF05739). When the SNARE database was used to analyse the proteins, a SNARE domain was identified (residues 259-312 in BCVIC2-1 with an e-value of 2.8e-11 and 226-279 in BCVIC2-2, with an e value of 3.1e-11) that in both proteins was classified as belonging to the Qa.IV subgroup and therefore putatively involved in exocytosis at the plasma membrane. The *Bcvic2-1-* and *2-2*-encoded amino acid sequences shared low identity, with a percentage identity of 65.9% over the entire sequence. The N-terminal regions from nucleotide position 1 to 201 only shared a pairwise identity of 33.3% and the sequences downstream towards the C terminus were more conserved, with a pairwise identity of 95.4%.

### All gene knockout transformants are initially heterokaryotic

Three independent transformation experiments using a combination of circular plasmid and linear split marker PCR fragments targeting *Bcvic1* and *Bcvic2* were required to obtain sufficient hygromycin-resistant transformants for further molecular analysis. For each gene transformants, were selected for downstream analysis. There was high efficiency of targeted integration, with only 13% of recombinants displaying ectopic integration of the gene knockout constructs. Diagnostic PCR indicated that there were some transformants with integration of a single flank at the homologous locus. Both these and any ectopic transformants were removed from the downstream analyses. However, all transformants with homologous integration of the knockout construct at both flanks at the desired locus were heterokaryotic, with diagnostic PCR indicating retention of the wild-type allele for the targeted gene (S3 Fig.). In the generation of control strains, the transformation efficiency of the non-targeted hygromycin resistance ectopic expression construct was markedly low in contrast to the targeted gene knockout constructs. Nevertheless, three independent mitotically stable hygromycin-resistant 839-5 and 839-1 nit tester isolates were selected for downstream analyses (S1 Table).

To generate homokaryon knockout lines, the heterokaryotic knockout mutants were purified by single-spore isolation. Conidia that germinated and grew robustly on MEA+Hyg100 were typically favoured to select against heterokaryotic conidia containing a large proportion of hygromycin-sensitive wild-type nuclei.

One round of single-spore isolation was sufficient to purify homokaryotic *ΔBcvic1* mutants. The majority of the single-spore isolates were homokaryotic: only 11 out of 75 tested positive for the presence of *Bcvic1*, as indicated by a faint PCR amplification product on an agarose gel. To ensure homokaryosis, a subsequent round of single spore isolations (*n*=15) was conducted and PCR analysis indicated absence of the *Bcvic1* wild-type allele in all the second round single-spore isolates.

In contrast to the ease with which homokaryotic *ΔBcvic1* mutants were obtained, the isolation of homokaryotic *ΔBcvic2* and *ΔΔBcvic1/2* mutants proved much more elusive. Many of the germinating *ΔBcvic2* and *ΔΔBcvic1/2* mutants were slower-growing germlings than the *ΔBcvic1* mutants. Nevertheless, 25–50 of the relatively faster-growing germlings were isolated and molecularly analysed based on the success of this strategy for the isolation of *ΔBcvic1* homokaryotic transformants. However, diagnostic PCR analysis of the viable isolates showed that all the *ΔBcvic2* and *ΔΔBcvic1/2* single-spore transformants were heterokaryotic since they were positive for the presence of the native gene (data not shown).

Despite four rounds of sequential single-spore isolations, no *ΔBcvic2* and *ΔΔBcvic1/2* homokaryotic transformants were isolated, suggesting that the *Bcvic2* gene might be essential or required for normal growth. If the *Bcvic2* gene was indeed essential or required for normal growth, it was assumed that the homokaryotic hygromycin-resistant mutants would either be nonviable (lethal phenotype) or grow abnormally slower than the heterokaryotic germlings. Based on this assumption, single-spore isolations were made from the heterokaryotic *ΔBcvic2* and heterokaryotic *ΔΔBcvic1/2* cultures, favouring slow-growing germlings with abnormal morphologies. One round of single sporing resulted in the isolation of homokaryotic *ΔBcvic2* or *ΔΔBcvic1/2* (confirmed by diagnostic PCR; S4 Fig.).

### Deletion of *Bcvic1* alone does not affect vegetative compatibility

Vegetative incompatibility tests were performed on three independent homokaryotic *ΔBcvic1* mutants (Fig. 4; S1 Table). *ΔBcvic1-i* and *ΔBcvic1-ii* were knockout mutants in the 839-5 *nit1* background, whereas *ΔBcvic1-iii* was created in the 839-5 NitM background. The nit mutant complementation tests showed that none of the *ΔBcvic1* mutants displayed any change in VI phenotype compared with the background 839-5 strain. All the mutants displayed vegetative compatibility with 839-5 and incompatibility when paired with 839-1. Three replications of the complementation experiments confirmed that the heterokaryon formation ability of the *ΔBcvic1* mutants was identical to that of the background 839-5 strain, indicating that the deletion of *ΔBcvic1* alone had no effect on vegetative incompatibility.

**Fig. 4.**
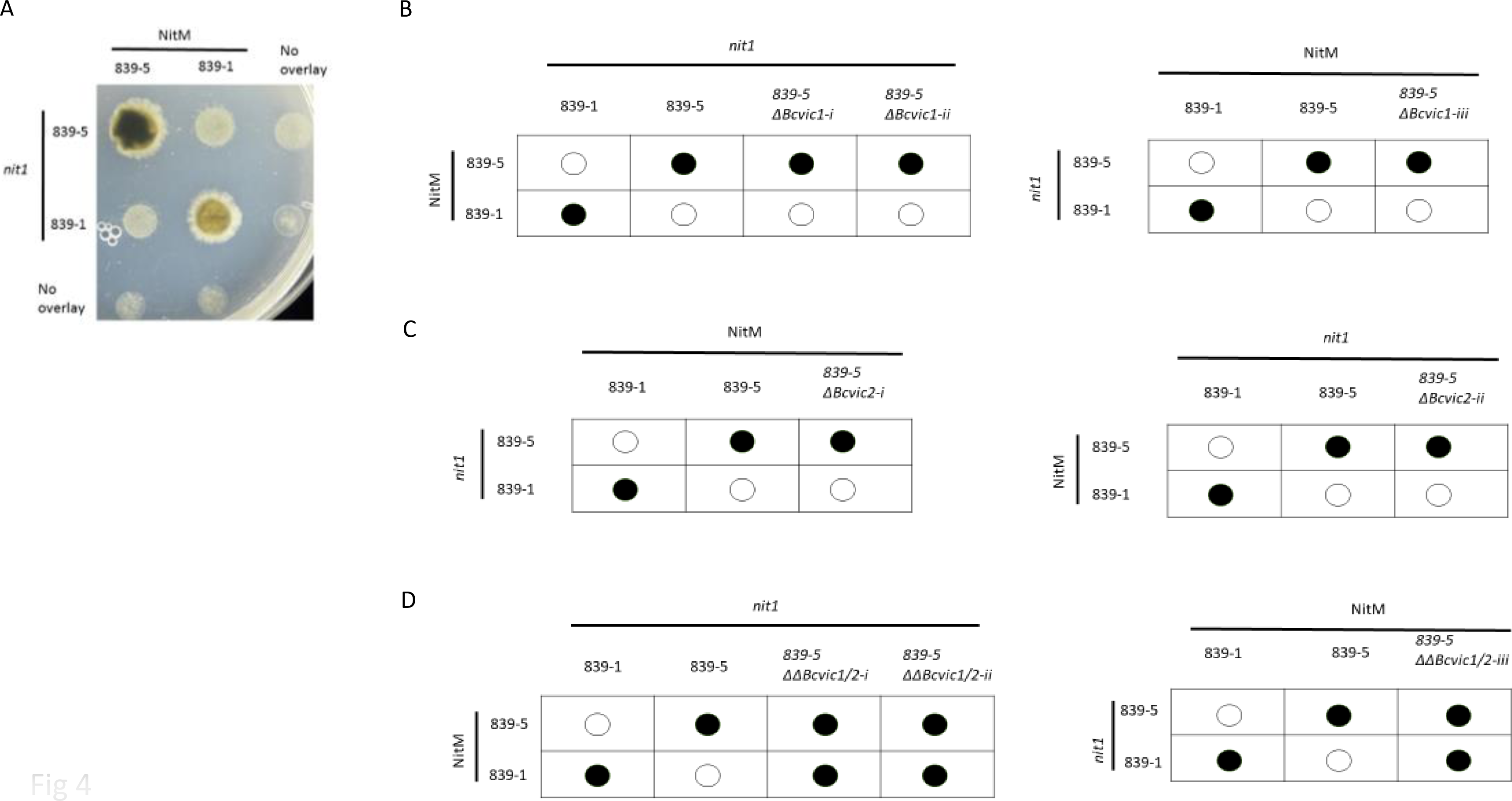
Nit mutant complementation tests for vegetative compatibility. (A) Complementation tests between *Botrytis cinerea* 839-5 and 839-1 *nit1* and NitM mutants. The overlaid Nit1 and NitM mutants were inoculated as drops of spore suspensions on MM+NO3 + triton and incubated for 10 days. Sparse growth indicates vegetative incompatibility, whereas dense growth indicates vegetative compatibility. (B-D) Schematic of nit mutant complementation vegetative compatibility tests for the *ΔBcvic1* mutants (B), the *ΔBcvic2* mutants (C) and the *ΔΔBcvic1/2* double knockout mutants (D). The black and white circles represent complementation and no complementation, respectively.

### *Bcvic2* is required for normal growth rate and habit

Deletion of the *Bcvic2* gene had a major morphological effect on fungal colony formation. At 30 h post-inoculation, transformants with an intact *Bcvic2* gene had elongated and branching hyphae that had grown ten times the length of hyphae in the *ΔBcvic2* or *ΔΔBcvic1/2* transformants, which had an atypical dwarf-like appearance, with engorged, shorter intertwined branches (Fig. 5). At 3 days post-inoculation, severe dysfunction in apical extension of the hyphae was evident, with an increase in apical and lateral branching that gave the colony a shortened fan-like shape of only approximately 1 mm diameter. This was in stark contrast to the typically fine, white mycelium of the heterokaryotic mutant or wild-type, which spread over a third of a 9-cm Petri dish. At 5 days post-inoculation, homokaryotic colonies appeared dark, with the formation of bulging conidiophore-like structures. At 14 days post-inoculation, the heterokaryotic mutant or wild-type isolates had colonised the entire plate with profuse sporulating hyphae, in contrast to the constricted homokaryotic colonies which were restricted to approximately 4 mm diameter, which by 60 d post-inoculation had only extended to a diameter of 35 mm with sclerotial bodies (Fig. 5). There were also long, hairy conidiophore-like structures protruding apically that were fragile and easily dislodged with light manipulation. These abnormal morphological characteristics were conserved in both *ΔBcvic2* and *ΔΔBcvic1/2* mutants (Fig. 5).

**Fig. 5.**
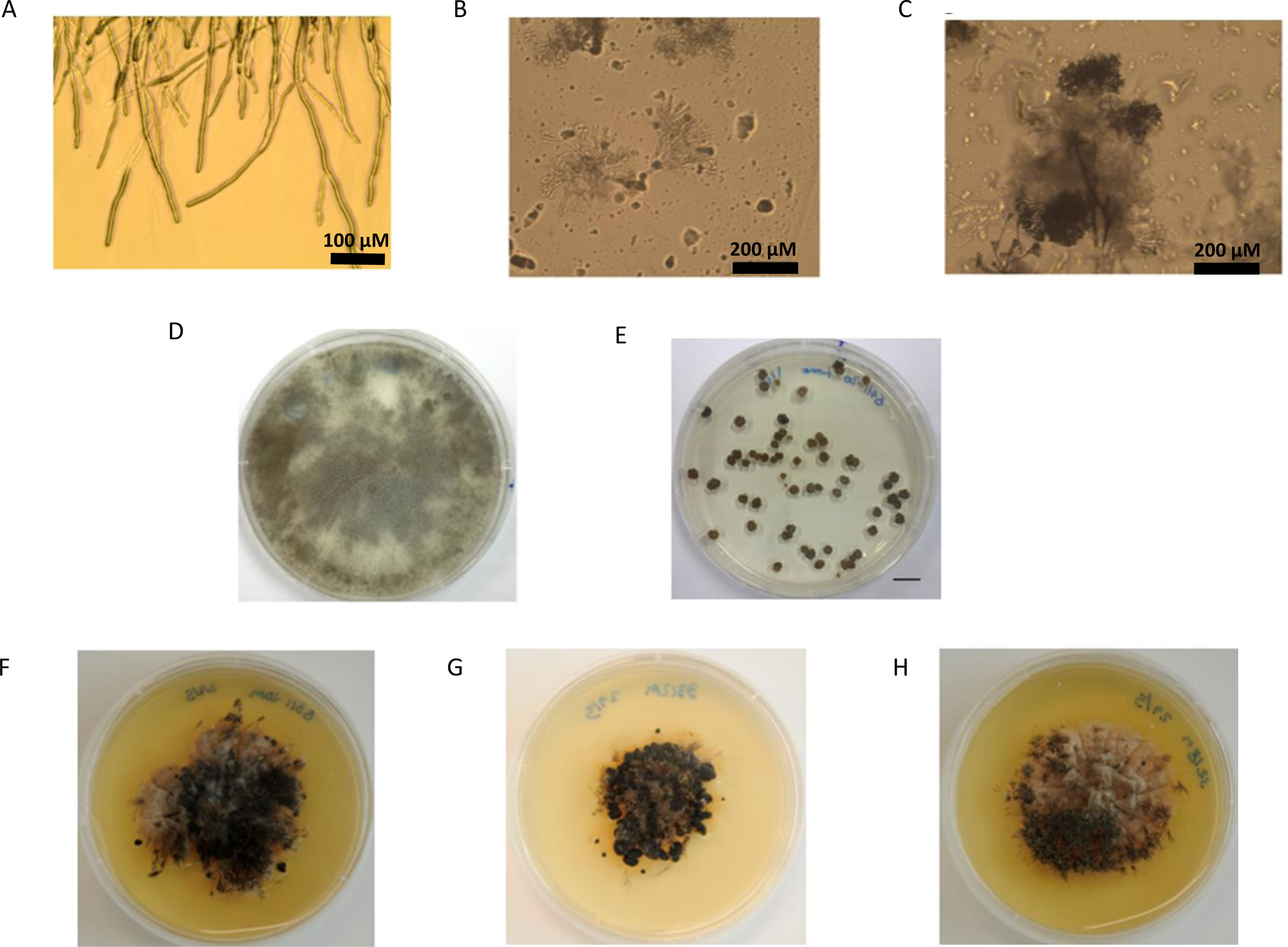
Abnormal hyphal growth and colony formation of the homokaryotic *ΔBcvic2* and *ΔΔBcvic1/2* mutants. (A, B) *HetΔΔBcvic1/2* and *ΔΔBcvic1/2*, respectively, at 3 days post- inoculation. (C) *ΔΔBcvic1/2*, at 5 days post-inoculation. (D, E) Spread plates of *HetΔΔBcvic1/2* and *ΔΔBcvic1/2*, respectively, at 14 days post-inoculation. (F, G) Two separate *ΔΔBcvic1/2* mutants at 60 days post-inoculation. (H) *ΔBcvic2* mutant at 60 days post-inoculation.

Initial problems were encountered during attempts to complement *ΔBcvic2* and *ΔΔBcvic1/2* mutants with wild-type alleles of the corresponding genes: the transformants were recalcitrant to protoplast isolation. To circumvent this problem, a secondary incubation step was introduced to generate young mycelial material following maceration of mature melanised hyphae (S5 Fig.). Transformation efficiencies equivalent to wild-type levels were achieved only by extending the incubation period for protoplast regeneration from 16 hours to 30 hours.

When the *ΔBcvic2* mutant was transformed with the *Bcvic2-1* ectopic expression construct, the growth phenotype of the *ΔBcvic2* + *Bcvic2-1* complementation mutant was similar to that of wild-type 839-5. These complementation results confirm that deletion of *Bcvic2* is indeed the cause of the abnormal growth phenotype in the *ΔBcvic2* mutant. The *ΔΔBcvic1/2* mutant was transformed with the *Bcvic1-1*, *Bcvic1-2*, *Bcvic2-1*, *Bcvic1-1/Bcvic2-1* and *Bcvic1-2/Bcvic2-2* complementation constructs (S1 Table). Complementation with *Bcvic1-1* and *Bcvic1-2* did not restore the colony morphology to wild type, whereas transformation with *Bcvic2-1*, *Bcvic1-1/Bcvic2-1* and *Bcvic1-2/Bcvic2-2* resulted in fast-growing colonies similar to the wild type (S6 Fig.). These results further confirm that deletion of *Bcvic2* and not *Bcvic1* results in the abnormal growth phenotype, as complementation with either *Bcvic1-1* or *Bcvic1-2* alone did not restore the colony morphology to the wild type.

### Deletion of both *Bcvic1* and *Bcvic2* is required to abolish vegetative incompatibility

*ΔBcvic2* mutants displayed vegetative compatibility phenotypes identical to that of the wild type; and compatibility with 839-5 and incompatibility when paired with 839-1, demonstrating that the deletion of *Bcvic2* alone does not affect vegetative incompatibility. In contrast, the deletion of both *Bcvic1* and *Bcvic2* resulted in the abolition of vegetative incompatibility, with mutants compatible with both 839-5 and 839-1 (Fig. 4).

Complementation of *ΔΔBcvic1/2* with *Bcvic1-1/Bcvic2-1* alleles restored vegetative incompatibility with 839-1 (Fig. 6). This result confirms that deletion of both *Bcvic1-1* and *Bcvic2-1* is required for the loss of the incompatibility phenotype. Interestingly, complementation of the *ΔΔBcvic1/2* mutant with *Bcvic1-2/Bcvic2-2* alleles from the previously incompatible tester 839-1 resulted in vegetative compatibility with 839-1 and incompatibility with the previously compatible 839-5. This reversal of vegetative compatibility phenotype indicates that interaction with 839-1 and 839-5 is allele-specific and further confirms that deletion of both *Bcvic1* and *Bcvic2* is required for the loss of the incompatibility phenotype in the *ΔΔBcvic1/2* mutant.

**Fig. 6.**
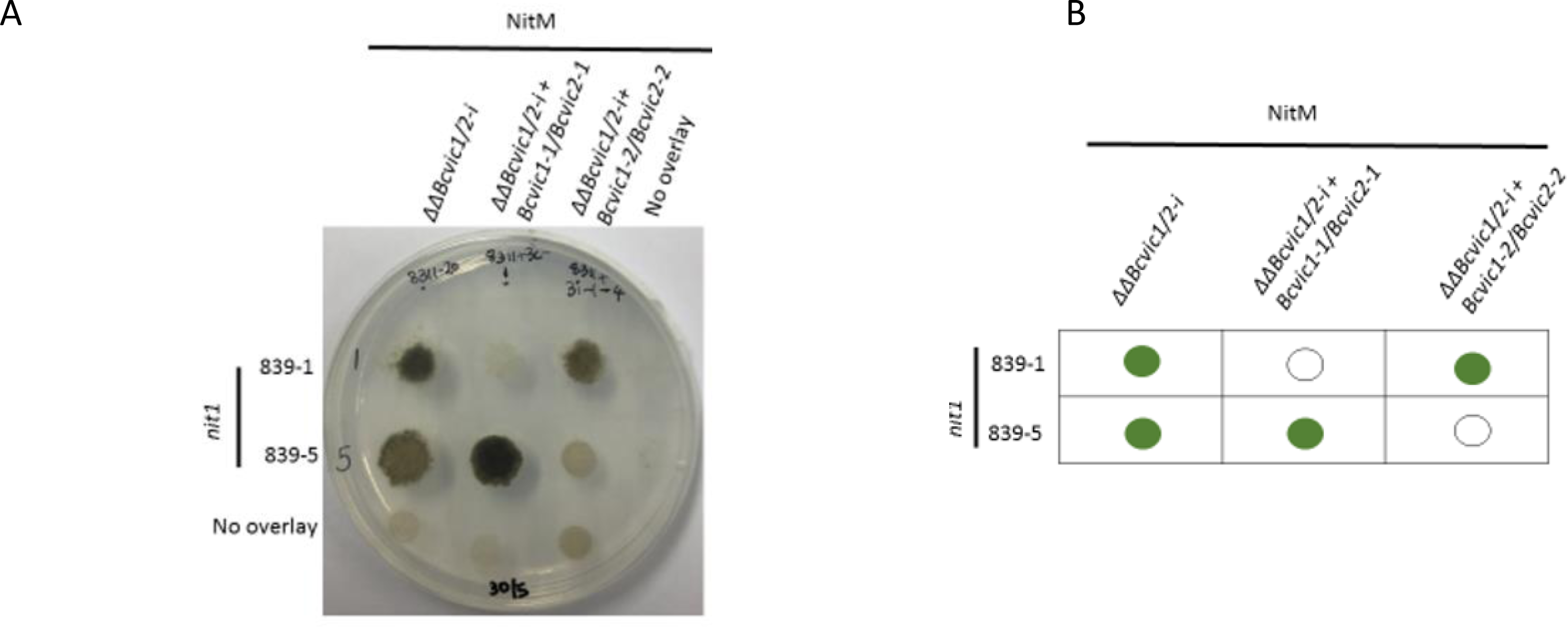
Complementation of *ΔΔBcvic1/2* with the *Bcivc1-1-Bcvic2-1* and *Bcviv1-2- Bcvic2-2* alleles. Nit mutant complementation to test vegetative compatibility between *Botrytis cinerea* 839-5 and 839-1 *nit1* and NitM mutants tester strains. The overlaid *nit1* and NitM mutants were inoculated as drops of spore suspensions on MM+NO3 + triton and incubated for 10 days. Sparse growth (white circle) indicates vegetative incompatibility, whereas dense growth (green circle) indicates vegetative compatibility. (A,B) Image and schematic of agar plate, respectively.

## Discussion

The molecular characterisation of allorecognition in filamentous fungi has relied on the dissection of the phenomenon in a limited number of species: *N. crassa, P. anserina and B. parasitica* [16]. With the characterisation of the first *vic* genes in *B. cinerea*. this report has expanded the number of fungi investigated The research strategy chosen greatly facilitated the identification of these genes. Genome-wide association studies (GWAS), including bulked segregant analysis (BSA) combined with whole genome/transcriptome sequencing, where the bulks are derived from progeny from a cross, have not been widely utilised in identifying genes of interest in fungi. However, these strategies have proved successful in identifying genes encoding virulence factors and candidate effectors from a number of species (e.g. [55–57], including genes involved in allorecognition in *N. crassa* [13]. In *B. cinerea* NGS-BSA has been used recently to identify a major-effect gene controlling pathogenicity and development [58]. The use of a backcrossed population for BSA significantly strengthens the strategy. The length of time taken to produce near-isogenic lines of *B. cinerea* may be viewed as a limiting factor, as each sexual cycle can take four to six months. However, for the purpose of identifying and testing *vic* genes, the backcrossing procedure was invaluable, since a smaller population sample size could be used for the mapping studies. In addition, the generation of nit mutants as a tool for determining the VI phenotype proved to be beneficial, as barrage tests, reliant on a morphological change (often characterised by pigmented, aerial mycelial growth) where incompatible hyphae meet can give somewhat ambiguous results [59]. Furthermore, the use of the nit mutant complementation assay provided a rapid VI phenotype result compared with the barrage test. However, an important consideration when choosing a VI assay is that nit mutant complementation assesses for the ability of two strains to form a heterokaryon, whereas the barrage test is a good measure of cell death after an incompatible reaction.

Identification of the two VI genes in *B. cinerea* has revealed both commonalities with, and differences from, genes involved in allorecognition in the other previously well-studied species. The two *B. cinerea vic* genes, *Bcvic1* and *Bcvic2*, have alleles which are highly polymorphic. This is a common characteristic of previously identified *vic* genes in other fungi, such as *het-c* in *N. crassa* and *het-D* in *P. anserina* [20, 34]. Furthermore, both the *Bcvic* genes share similarity to allorecognition genes in other fungi.

The *Bcvic1* gene encodes an NLR protein with a tri-partite domain structure. NLRs have been identified in numerous allorecognition systems in fungi [13, 30, 34, 60, 61], but whilst the overall NLR architecture of BCVIC1 is typical of members of the receptor family, both common and novel domains are present. The central NACHT domain, which has a nucleotide-binding site, is shared by a number of VI proteins in the three well-studied species [16]. In contrast, C-terminal ankyrin repeat domains have not been previously attributed to VI proteins, but alternatives which putatively share similar functionality (probably driving protein-protein interactions), most notably WD40 repeat (WDR) or tetratricopeptide (TPR) repeat domains, predominate [13, 30, 34, 60, 61]. The N-terminal putative lipase domain is common to a number of genes involved in allorecognition [13, 60]. *Bcvic2* encodes a putative syntaxin protein with a SNARE domain which has a predicted function in vesicular transport; similar proteins are found as components of the VI systems of *N. crassa* and *P. anserina* (*sec-9*; [13]), as well as *C. parasitica* (*vic2a*; [60]). Thus, the first VI system characterised in *B. cinerea* appears to be analogous to the systems in other fungi that rely on NLR/SNARE proteins.

Of these systems that rely on NLR/SNARE proteins, the *Plp-1*/*sec-9* system of germling-related death (GRD) in *N. crassa* is probably the best characterised [13]. PLP-1 is a typical tri-partite NLR, which physically interacts with SEC-9, a SNARE-domain protein, of differing GRD specificity, to initiate the VI response. The specific recognition interaction is thought to involve the C-terminal TPR domain of PLP-1, with a functional Nucleotide-Binding Domain (NBD), the central NB-ARC domain, required for oligomerisation of PLP-1 following activation. A functional N-terminal patatin-like phospholipase domain is required for translation of recognition into the cell death response, with NB-ARC domain-mediated self-association being insufficient to result in GRD. Owing to the similarity of domain architecture, it is highly likely that the *Bcvic1*/*2* system in *B. cinerea* operates in a similar manner. In BCVIC1, the ankyrin repeats probably mimic the TPR domain of PLP-1 in their functionality, as both can act as scaffolds to mediate protein–protein interactions [62, 63], with BCVIC1 perhaps physically interacting with BCVIC2, of differing specificity, leading to PCD. The NACHT domain, predicted to have nucleoside-triphosphatase (NTPase) activity, is functionally equivalent to the NB-ARC domain of PLP-1, and potentially required for oligomerisation. The N-terminal putative lipase domain of BCVIC1 is probably required for the cell death response; however, despite advances in identifying the genes and encoded proteins involved in fungal VI, how the PCD response is effected is poorly understood.

Deleterious alteration of membranes is a common early event in the initiation of many VI-associated cell death responses [16]. Uncovering the functionality of proteins involved in orchestrating VI has revealed how membrane perturbation is effected during PCD initiation. For example, the *regulator of cell death* (*rcd-1*) gene involved in VI in *N. crassa* resembles the N-terminal domain of gasdermin, which forms pores in mammalian cell membranes that disrupt trans-membrane ion gradients leading to pyroptotic cell death, a key component of mammalian innate immunity. Coexpression of incompatible *rcd-1* and *rcd-2* alleles triggers pyroptotic-like cell death in human 293T cells [14, 64]. The postulated functionality of a number of VI-associated NLR N-terminal domains also suggests how membranes may be compromised to initiate PCD. The enzymatic activity of the patatin-like phospholipase domain of PLP-1 is essential for PCD, with this activity potentially able to compromise membrane phospholipids directly, leading to cell death. This activity could be shared by both the putative lipase domain in BCVIC1 and the patatin domain of VIC2 from *C. parasitica* [60]. In contrast to PLP-1, the predicted PCD-initiating function of the N-terminal domain of BCVIC1 may not rely on enzymatic activity. Fungal NLRs share architecture with NLRs involved in animal and plant immunity [65–67]. Solving the structure of animal and plant NLRs has enabled functionality to be analysed, as exemplified by ZAR1, the first plant NLR to have its structure solved [68, 69]. When ZAR1 recognises its cognate bacterial virulence protein, it forms a pentameric structure on activation, termed a resistosome [68, 69]. The activated ZAR1 resistosome acts as a calcium-permeable cation channel, with the influx of Ca^2+^ leading to immunity and cell death, resulting from perturbation of subcellular structures and the generation of ROS [70]. The initiation of PCD by BCVIC-1 following recognition of BCVIC-2 may also rely on the formation of a cation channel. Solving the structure of BCVIC1 and BCVIC2 may shed light on this possibility.

BCVIC2, because of its conserved syntaxin and SNARE domains, and its putative classification as a Qa.IV SNARE protein, is probably involved in exocytosis at the plasma membrane [71–73], and was shown to be essential for normal vegetative growth. Indeed, in other pathogenic filamentous fungi, SNARE proteins are involved in asexual development, stress tolerance and pathogenicity [74–76]. Heller and colleagues [13] postulated that the SEC- 9 SNARE-domain proteins in *N. crassa*, *P. anserina* and *C. parasitica* are targeted by effectors from other microbes that aim to inactivate exocytosis/autophagy, thereby driving intraspecific polymorphism. This has led to their recruitment in innate immunity as the guardees in an NLR- guarded system, with exaptation resulting in this system then being exploited in allorecognition with the repeated recruitment of the PLP-1/SEC-9 pair. Indeed, BCVI2 is predicted to belong to the Qa.IV subset of SNARE proteins involved in secretion, and these proteins are putatively more highly variable than the conserved proteins involved in membrane fusion events at the endoplasmic reticulum and Golgi apparatus [77]. Although BCVIC-1 and 2 only share domains and overall predicted structure/function with PLP-1/SEC-9 of *N. crassa*, *B. cinerea* is the fourth fungus in which an NLR-SNARE domain protein pair has been demonstrated to function in allorecognition. Furthermore, *Botrytis* is the first example from a fungal species that is not a Sordariomycete, lending further support to the generality of these hypotheses. Specific modifications of a guarded SNARE protein that leads to the NLR initiating a cell death response have not been investigated in any of the fungal VI systems characterised to date. However, since BCVIC2 is putatively designated to act at the plasma membrane, its activation, promoted by Sec1/Munc18 (SM) regulatory proteins or posttranslational modifications, to enable it to participate in forming a SNARE complex [78], or its interaction with tethering factors involved in targeting of membrane fusion events [79], may be sufficient to trigger the guarding NLR.

In previous VI studies in other fungal systems, rescue of homokaryotic knock-out mutation lines of the gene encoding the SNARE-domain protein proved impossible [13, 52]. Similarly, isolating homokaryotic lines of *ΔBcvic2* and *ΔΔBcvic1/2* mutants from heterokaryotic cultures initially proved difficult. This suggested that *Bcvic2* may be essential, paralleling the previous results from *C. parasitica*, *P. anserina* and *N. crassa* [13, 52]. However, the *ΔBcvic2* and *ΔΔBcvic1/2* homokaryon deletion lines were finally obtained after the single-spore isolation strategy was altered to select for abnormal phenotypes, with slow growth. This finding highlights the importance of removing bias when selecting single-spore isolates. Furthermore, it also highlights the importance of retaining isolates that appear non-viable, and using microscope analysis to confirm viability versus non-viability, when phenotype may be equivocal. These considerations are made apparent by Ko and colleagues [80], in which the authors were able to demonstrate the function of a gene essential for cell division and polarised growth, cell division cycle 48 (CDC48), in *C. parasitica*. On initial inspection, no growth of spores from a heterokaryotic transformant on selective media was observable, suggesting that *Cpcdc48* was essential. However, extended incubation combined with microscopic analysis revealed transient, aberrant germination of some conidia prior to autolysis, indicating compromised cell division and polarised growth [80].

Identification of the first genes involved in allorecognition in *B. cinerea* will enable further studies on the mechanisms governing PCD in this fungus. An understanding of these intrinsic cell death pathways, the components and the signalling cascades involved will provide targets for manipulation, for example, using spray-induced gene silencing to alter expression of genes encoding key elements [81, 82], so that PCD can be prematurely initiated to prevent disease. Uncovering further genes involved in allorecognition in *B. cinerea* may also pave the way for the development of a super donor strain capable of transmitting hypovirulent mycoviruses that lack a mechanical transmission route [49–51] like that developed for *C. parasitica* [52, 53]. Whilst a mechanically transmissible hypovirulent virus has been discovered in *B. cinerea* [83], the necessity of multiple applications for disease control may reduce the practicalities of its use, thus, development of such a donor would enable exploitation of hypovirulent mycoviruses as potent BCAs against *B. cinerea*.

## Methods

### Fungal strains

#### Culturing

*B. cinerea* wild-type strains were routinely cultured on malt extract agar (MEA, Oxoid, Basingstoke, UK) or potato dextrose agar (PDA, Difco™, BD Biosciences, Franklin Lakes, NJ, USA) in darkness. To induce sporulation, 3- to 4-day-old cultures were exposed to near-UV light (350–400 nm) for 16 hours, and were subsequently returned to darkness. Conidia were harvested 4–7 days later. Sporulating cultures were flooded with sterile reverse osmosis water (SW) and spores dislodged to prepare spore suspensions. Following centrifugation at 11,000 *g* for 5 minutes spore suspensions were adjusted, if required, to the desired concentration in SW. Isolates and strains used in this study are listed in S1 Table.

#### Mating type

Mating type was established by diagnostic routine PCR (see section below) to enable selection of isolates for subsequent backcrossing. Mating type-specific primers were designed to differentiate between the *MAT1* and *MAT2* loci (S3 Table). *MAT1*-specific primers (MAT1F/R) were designed against the 5’ portion of the *MAT1-1-1* coding sequence (CDS) retrieved from the B05.10 v1 genome sequence (downloaded from the Broad institute www.broadinstitute.org/annotation/genome/botrytis_cinerea [47], now located on the Joint Genome Institute MycoCosm site (https://mycocosm.jgi.doe.gov/Botci1/Botci1.home.html; GeneID: BC1G_15148). *MAT2* specific primers (MAT2F/R) were designed against the *MAT1- 2-1* CDS retrieved from the T4 genome sequence ((https://urgi.versailles.inra.fr/Species/Botrytis) ([84]; GeneID: BofuT4_P160320.1).

#### Antibiotic resistance

Strains were tested for benzimidazole fungicide resistance by plating 10 µL of a dense spore suspension on to MEA + vinclozolin (100 mg/L) and assessing growth at 3 days, with resistant strains producing a compact mycelial mat. For dicarboximide resistance, the spore suspension was plated onto MEA + carbendazim (100 mg/L) and assessed at 2 days, with resistant strains producing spreading colonies.

#### Mycelial compatibility

Generation of nitrate non-utilising mutants to demonstrate mycelial compatibility resulting in heterokaryon formation followed the method of Beever and Parkes [59]. For each strain, 3-mm mycelial plugs from 3-day-old MEA cultures were transferred mycelial side down on Vogel’s N- minimal medium [85] amended with potassium chlorate (30 g/L; MM+ ClO3) and incubated at 20°C in the dark for up to four weeks. Sectors arising at colony margins were purified by transferring mycelial plugs to fresh MM+ ClO3. Mycelial plugs from the colony margin of the sub-culture were then transferred onto MEA plates for growth and induced to sporulate. Conidia were harvested after 7 days and stored as water cultures for further phenotype testing.

Putative *nit1* (able to utilise nitrite, ammonium, hypoxanthine and uric acid) and NitM (able to utilise nitrite, ammonium and uric acid, but not hypoxanthine) mutants were classified by plating mycelial plugs onto MEA amended with four nitrogen sources and uric acid following the procedure of Beever and Parkes (2003). Growth was scored after 6–7 days for either wild type growth (+) or ‘minus nitrogen’ (-) sparse growth similar to growth on nitrogen-free medium. Complementation assays to test for vegetative compatibility were performed by overlaying spore suspensions of *nit1* and NitM mutants of the isolates being tested on Vogel’s N- [85] + NO3 medium amended with triton X-100 (0.5 mL/L) followed by incubation for 8–14 days at 20°C. A dense mycelial pad indicated successful heterokaryon formation, whereas absence of complementation was indicated by sparse growth.

### Fungal genomic DNA extraction

For preparing genomic DNA for Illumina sequencing and as a template for confirmation of mating type and candidate gene sequences, fungal material was generated by inoculating 100 mL of potato dextrose broth (PDB; Difco™, BD Biosciences) with spore suspensions derived from sporulating cultures, which were then incubated statically for 30 hours to reduce polysaccharide production. Germlings were harvested by centrifugation at 8,500 *g* for 15 min at 4°C (Sorvall™ RC6 Plus, Thermo Fisher Scientific, Waltham, MA, USA), washed once with SW and re-centrifuged prior to being used as the starting fungal material for genomic DNA extractions. Fungal DNA extractions were performed according to the method of Möller et al. [86]. DNA was extracted from 1.2 g of fresh fungal material and finally resuspended in 200 µL of SW.

For rapid screening of transformants, genomic DNA was extracted from a small amount of mycelia and conidia (30–60 mg) scraped from a plate of fungal culture. This material was placed into a 1.5-mL safe-lock tube (Eppendorf, Hamburg, Germany) containing three 3.2-mm stainless steel beads (Next Advance, Troy, NY, USA) and 500 µL of TES (100 mM Tris, pH 8.0, 10 mM EDTA, 2% SDS). The fungal material was then macerated using a Bullet Blender tissue homogeniser (Next Advance) for three 1 min cycles at speed level 12. After homogenisation, extractions were performed according to a scaled-down version of the protocol of Möller et al. [86], with the purified DNA being finally resuspended in 50 µL of dH20. DNA concentrations were quantified using the NanoDrop™ 2000 spectrophotometer (Thermo Fisher Scientific, Waltham, MA USA) and integrity verified by gel electrophoresis.

### Polymerase chain reaction (PCR)

All PCRs were carried out using a Mastercycler Gradient machine (Eppendorf) and amplification products were visualised by gel electrophoresis. Routine PCR was used to amplify target DNA using platinum *Taq* DNA polymerase (Invitrogen) according to the manufacturer’s instructions. PCR fragments that were destined to be cloned or sequenced directly were amplified using the Q5 high-fidelity DNA polymerase (New England Biolabs, Ipswich, MA, USA) according to the manufacturer’s instructions.

### Bulked segregant analysis

#### Isolation of near-isogenic lines by backcrossing

Near-isogenic strains of *B. cinerea* were generated by a backcrossing programme (S1 Fig.) with crosses performed as previously described [87, 88]. Initial parental strains, SAS405 (International Collection of Microorganisms from Plants (ICMP), Manaaki Whenua – Landcare Research, Auckland, New Zealand, ICMP10935) and REB704-1, differing in benzimidazole and dicarboximide sensitivity, were crossed to generate REB749-8 (ICMP15036). The SAS56 (ICMP10934) strain was then crossed with REB749-8 to generate the isolate REB800-4. This isolate was then used in the first of three backcrosses against the recurrent parent REB749-8. For the backcrosses, REB749-8 was used as the female sclerotial parent. At each generation, progeny that were incompatible with both parents were selected to fertilise the recurrent parent REB749-8. Five crosses were set up for each generation and only true crosses that had a 1:1 segregation of the fungicide resistance markers were selected for downstream experiments. The final F1BC3 population (n=32), was designated the REB839- series. REB839- single ascospore strains were either classified as *v*egetative *c*ompatibility *g*roup 1 (vcg1) or vcg2 by incompatibility testing against the recurrent parent (REB749-8) and the non-recurrent parent from F1BC2 (REB811-28) using nitrate non-utilising mutants, and bulked accordingly for the sequencing.

#### Illumina sequencing and read mapping

The DNA samples from each of the 32 REB839- series progeny were quantified using a NanoDrop™ 2000 spectrophotometer (Thermo Scientific™, Thermo Fisher Scientific), normalized to 100 ng/mL, and then pooled in equimolar amounts into a vcg1 bulk and a vcg2 bulk. Sequencing (100-bp paired end) of the two bulks was carried out in separate lanes on the Illumina Genome Analyzer II by the Australian Genome Research Facility (AGRF) following non-indexed library preparation.

The bulked vcg1 and vcg2 sequencing reads were trimmed to their longest contiguous region using the DynamicTrim and LengthSort modules in the SolexaQA package. Reads with Phred quality scores lower than an error probability of 0.05, and longer than 25 bp, but still paired, were retained for downstream analysis [89]. The trimmed vcg1 and vcg2 reads were then separately mapped to the unmasked reference genome of the *B. cinerea* T4 strain and the B05.10 strain (B05.10 v1), consisting of 118 and 588 scaffolds respectively, using the Burrows-Wheeler transform (BWT) algorithm implemented in the Burrows-Wheeler Aligner (BWA) programme v0.5.8 [90]. The resulting vcg1 and vcg2 alignment files were input into the Integrative Genomics Viewer (IGV) software for visualisation of the mapped reads [91].

#### Identification and analysis of single nucleotide polymorphisms (SNPs)

SNPs were identified relative to the T4 reference genome using a combination of tools embedded within the SAMtools package [92, 93]. To increase the fidelity of SNP calling, the polymorphism needed to i) be represented by at least eight reads; ii) occur at a position covered by at least eight reads in the other sequenced bulk; iii) not correspond to missing data in the reference strain; and iv) differ from the reference strain.

To identify outlier regions of the genome that contained an increased density of bulk-specific SNPs, a 5000-bp sliding window with a 25-bp lag was applied to the entire T4 genome sequence. The number of vcg1 and vcg2 bulk-specific SNPs was determined within each 5000-bp window. The 95^th^ percentile of the distribution was one SNP for both bulks, and therefore any region where both bulks showed more than one bulk-specific SNP within a 5000- bp window was considered a statistical outlier and a candidate location for the *vic* locus.

#### Identification of candidate vic genes

The complete sequence of the candidate *vic* region was obtained from the T4 sequence database. The predicted genes (called using the EuGene gene finding software [94]) within the candidate region in the T4 sequence were retrieved from the proprietary GnpGenome database embedded within the INRA website.

The candidate region was further analysed using a second gene prediction software, the FGENESH HMM-based gene structure prediction programme (www.softberry.com), using *B. cinerea*-specific gene-finding parameters to add additional evidence for the position of ORFs and intron/exon boundaries [95, 96]. Orthologous genes were identified in the candidate region of the gapless B05.10 genome [97] using BLASTn. The nucleotide and predicted amino acid sequences of the open reading frames shared between the two genomes, within the candidate region, were compared for the vcg1 and vcg2 bulks. The gene predictions were searched for the presence of domains or motifs using the Pfam software [98, 99], Interproscan 5 [100] and the SNARE database (http://bioinformatics.mpibpc.mpg.de/snare/index.jsp SNARE database [73]).

### Confirmation of candidate *vic* gene and predicted protein sequences

The sequences of the *Bcvic1* and *Bcvic2* candidate genes in the vcg1 and vcg2 tester strains were confirmed by PCR amplification, cloning and re-sequencing. Primers that spanned the *Bcvic1* and *Bcvic2* candidate genes, including the 5’ and 3’ UTR regions, were designed by manual inspection (S3 Table). The gene fragments were designed to overlap by 4 bp. The *Bcvic1* and *Bcvic2* primers used in the vcg2 tester, 839-1, were designed using the gapless B05.10 genome sequence, as they shared identical sequences. The vcg1 tester 839- 5 did not share sequence identity with B05.10 in the candidate region and therefore presented as a gap in the sequence. This sequence gap was cloned and resolved by primer walking by Macrogen Inc. (South Korea). Primers for the *Bcvic1* and *Bcvic2* genes in 839-5 were then designed based on that completed sequence. The coding sequence and predicted amino acid sequence of *Bcvic1* and *Bcvic2* in 839-5 (vcg1) and 839-1 (vcg2) were compared using a pairwise MUSCLE alignment in Geneious 8.1.2 (https://www.geneious.com).

### Functional characterisation of candidate genes by transformation

#### Construction of transformation vectors

For construction of fungal transformation vectors, required sequences were PCR-amplified, cloned individually into pCR XL-TOPO according to the manufacturer’s instructions, and then combined as required in the pType IIs vector via a Golden Gate strategy [101, 102]. To generate final transformation vectors, individual sequences from the pCR-XL-TOPO clones were assembled using the pType IIs vector (Invitrogen) as the backbone, according to the manufacturer’s instructions, using the *Aar*I typeII restriction enzyme for creation of non-palindromic overhangs, with the *Aar*I typeIIS restriction site engineered into all primers (S3 Table).

Plasmid DNA was extracted from 2 mL of *E. coli* bacterial overnight culture using the Zyppy plasmid miniprep kit (Zymo Research), or 200 mL of culture was processed with the PureLink HiPure pasmid midiprep kit (Invitrogen) according to the manufacturers’ instructions. To ensure correct assembly of clones, plasmid DNA was extracted and sequenced (Macrogen, South Korea) using either the M13F and M13R universal or gene-specific primers for pCR-XL-TOPO vectors or primers GGF and GGR for the pType IIs vector (S3 Table). Bioinformatic validation of sequences was conducted using Geneious 8.1.2 (https://www.geneious.com).

#### Cloning strategy for knockout transformation vectors

For knockout transformations, the hygromycin phosphotransferase resistance (*HPH*) gene was used as selectable marker, utilising a replacement strategy. Knockout vectors were constructed and split marker sequences engineered, composed of three separate fragments: a left flank, the HPH cassette, and a right flank. Primer sequences for generating the fragments for the knockout constructs are shown in S3 Table. The HPH cassette was PCR-amplified using the pNDH-OGG vector as the template [103] and the gene flanking sequences used 839-5 genomic DNA as the template. Whole vectors were used in downstream fungal transformations.

Pairs of split marker sequences were engineered with either 5’ or 3’ gene flank sequence and overlapping truncated fragments of the *HPH* cassette. Primer sequences are presented in S3 Table. The DNA templates for these reactions were the specific classical gene knockout vectors in the pType IIs backbone. The split marker fragments were used in downstream fungal transformations as purified PCR products.

#### Cloning strategy for complementation transformation vectors

Coding sequences of candidate *vic* genes and between 700 and 1000 bp of upstream and downstream sequences were PCR-amplified using genomic DNA from 839-5 and 839-1. The nourseothricin (*NAT*) selectable marker under the control of the *Aspergillus nidulans trpC* promoter was amplified from the pNAN-OCT vector [103]. The *A. nidulans trpC* (T:trpC) terminator was amplified from the pAN7-1 vector [104] and was used as the terminator for the *NAT* selectable marker. Whole vectors or linear PCR fragments encompassing the selectable marker and gene of interest, including the *A. nidulans* promoter and terminator, were used in downstream fungal transformations.

#### *B. cinerea* PEG-mediated protoplast transformation

##### Knockout transformations

The *B. cinerea* near isogenic vcg1 tester isolates, 839-5 *nit1* and 839-5 NitM, were used for PEG-mediated protoplast knockout transformation according to the procedure described by ten Have *et al.* (1998). Spore suspensions were used to inoculate 100 mL of PDB and allowed to germinate for 20 h in static liquid culture at 20°C. After incubation, the fungal material was harvested by centrifugation at 2500 *g* for 15 min and washed twice with KC buffer (0.6 M KCl and 50 mM CaCl2). The fungal material was then treated with either 10 mg/mL Glucanex (Sigma-Aldrich, St. Louis, MO, USA, discontinued) or 25 mg/mL lysing enzymes from *Trichoderma* (Sigma-Aldrich) to generate protoplasts which were then transformed with a combination of either 30 µg of circular plasmid DNA or 2.5 µg of split-marker PCR products. Transformed protoplasts were plated onto SH medium (0.6 M sucrose and 5 mM HEPES pH 6.5, 0.8% (w/v) agar) and allowed to regenerate for 24 hours. Transformation plates were then overlaid with 0.6% (w/v) water agar amended with 70 µg/mL hygromycin B (Invitrogen). Hygromycin-resistant colonies emerging on the surface of the overlay after 3–5 days were excised and transferred to MEA amended with 100 µg/mL of hygromycin B (Invitrogen) (MEA+Hyg100). To select for transformants that were mitotically stable, hyphal tips were excised and transferred to plates without selection and then subcultured back on to antibiotic The *B. cinerea* transformants were screened by PCR analysis. To confirm targeted left flank and right flank integration of the knockout constructs, primers were chosen outside the 5’ and 3’ flanks and within the HPH cassette. To confirm the presence or absence of the wild-type gene, primer pairs SA17/SA18 (384 bp) and SA19/SA20 (830 bp) were used and were specific to *Bcvic1* and *Bcvic2*, respectively. Specific primer sequences are listed in S3 Table.

##### Complementation transformations

The PEG-mediated transformation protocol [105] was amended for complementation transformations because of the slow growth rate and melanised nature of the knockout mutants. To maximize the fungal mass available for generating protoplasts, a loopful of crushed mycelia was spread over 90-mm Petri plates containing MEA+Hyg100 and incubated for 30 days after which melanised aerial structures were removed and crushed using a micropestle. The resulting slurry was used to inoculate 50 mL of PDB+Hyg100 and incubated statically for 5 days at room temperature and monitored daily for new hyphal growth. This fungal material was harvested and used as the starting material for the transformation.

Protoplasts were transformed with the complementation constructs, either 30–50 µg of the circular plasmid or 2.5 µg of the purified linear PCR product, and allowed to regenerate on SH media. To account for the slow-growing nature of the *ΔBcvic2* and *ΔΔBcvic1/2* mutants, the regeneration period was extended to 30 hours before overlaying with MEA amended with 100 µg/mL nourseothricin (MEA+NAT100). Nourseothricin-resistant transformants were excised and subcultured onto MEA+NAT100.

### Generation of homokaryon transformant lines

To obtain homokaryon lines from the heterokaryotic transformants, single-conidium isolations were performed. Conidia were serially diluted in SW and plated onto 0.6% (w/v) water agar supplemented with 100 µg/mL hygromycin B and incubated overnight at 20°C. Dilution plates that contained a sufficiently low density of conidia, as assessed under the light microscope, were used for isolations. Approximately 25 to 50 antibiotic-resistant germlings were excised and transferred onto MEA plates containing 100 µg/mL hygromycin B and allowed to grow at 20°C for 1 week and induced to sporulate. Mature conidia and mycelial fragments were harvested, DNA extracted and PCR used to confirm presence/absence of wild-type alleles (primers in S3 Table). If homokaryons were not purified after a single round of single-spore isolations, further rounds were completed until homogeneity was observed by PCR.

## Acknowledgments

We would like to thank Kim Snowden, Erik Rikkerink (both of The New Zealand Institute for Plant and Food Research Ltd) and Carl Mesarich (Massey University) for critically reviewing this manuscript.

## Author contributions

REB, MDT, MNP, SA and JKB conceived the experiments; SA, SLP and MPC performed the experiments; SA, MDT and JKB analysed the data; JKB, SA and MDT wrote the paper.

## Supporting information

**S1 Table.**
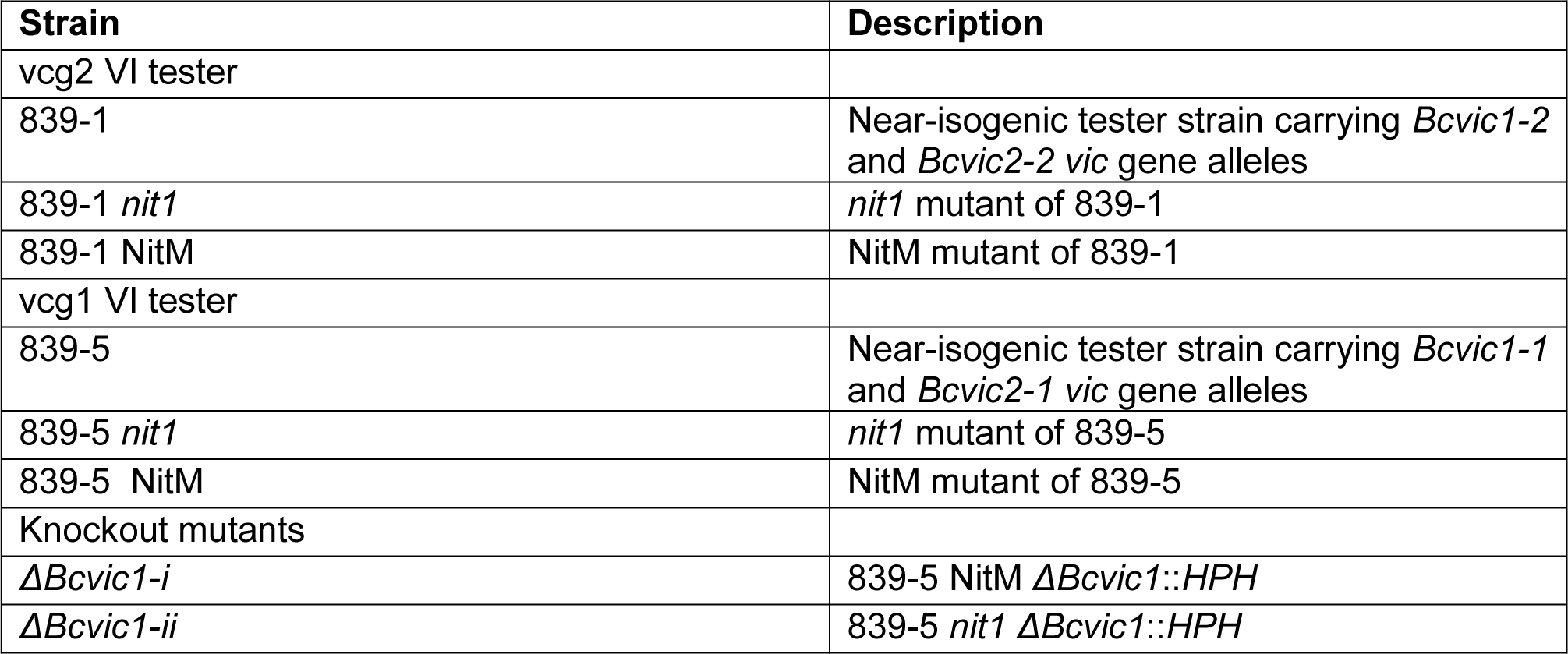
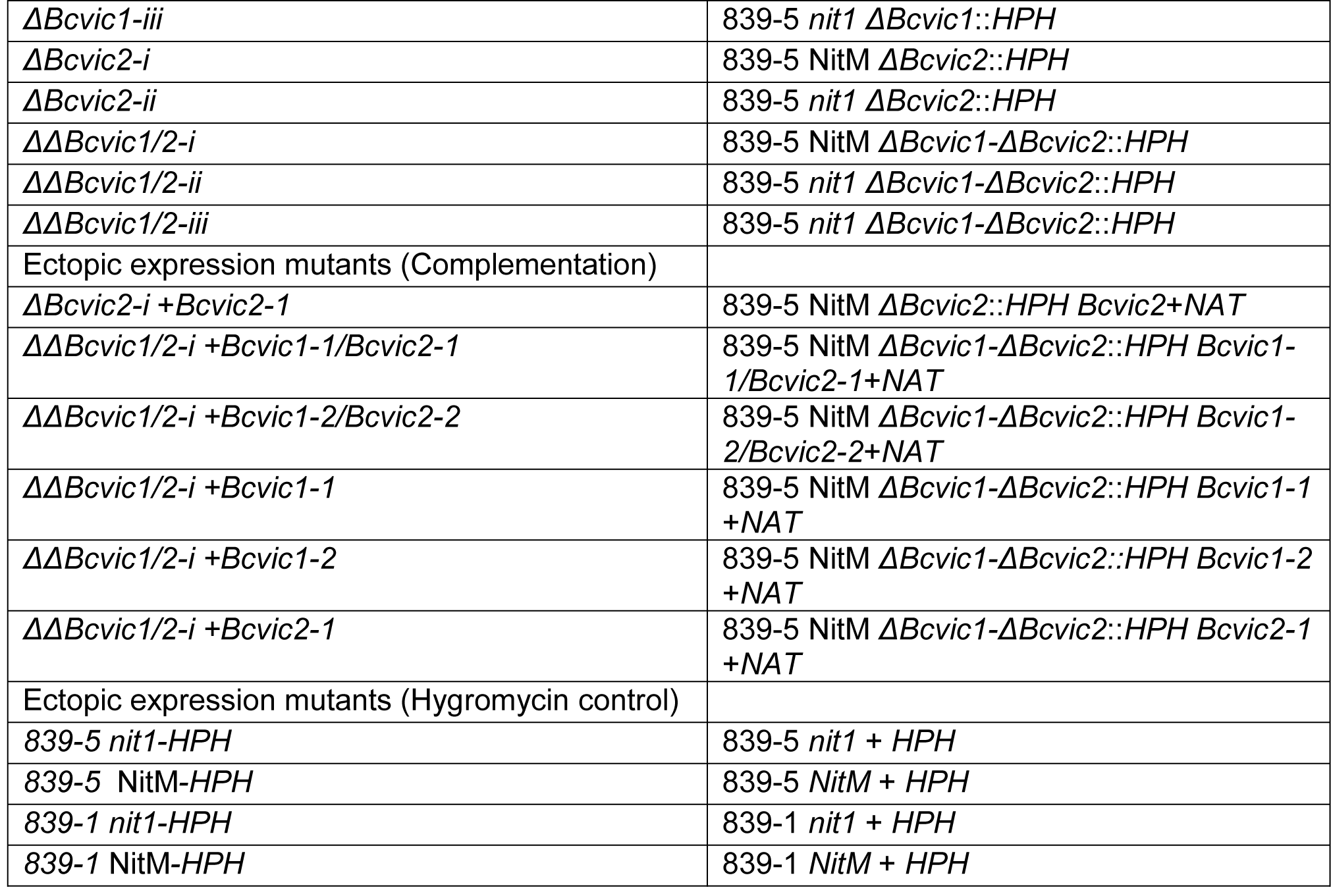
List of strains used in this study.

| Strain | Description |
| --- | --- |
| vcg2 VI tester |  |
| 839-1 | Near-isogenic tester strain carrying *Bcvic1-2* and *Bcvic2-2 vic* gene alleles |
| 839-1 *nit1* | *nit1* mutant of 839-1 |
| 839-1 NitM | NitM mutant of 839-1 |
| vcg1 VI tester |  |
| 839-5 | Near-isogenic tester strain carrying *Bcvic1-1* and *Bcvic2-1 vic* gene alleles |
| 839-5 *nit1* | *nit1* mutant of 839-5 |
| 839-5 NitM | NitM mutant of 839-5 |
| Knockout mutants |  |
| $\Delta Bcvic1-i$ | 839-5 NitM $\Delta Bcvic1::HPH$ |
| $\Delta Bcvic1-ii$ | 839-5 *nit1* $\Delta Bcvic1::HPH$ |
| *ΔBcvic1-iii* | 839-5 *nit1 ΔBcvic1::HPH* |
| *ΔBcvic2-i* | 839-5 NitM *ΔBcvic2::HPH* |
| *ΔBcvic2-ii* | 839-5 *nit1 ΔBcvic2::HPH* |
| *ΔΔBcvic1/2-i* | 839-5 NitM *ΔBcvic1-ΔBcvic2::HPH* |
| *ΔΔBcvic1/2-ii* | 839-5 *nit1 ΔBcvic1-ΔBcvic2::HPH* |
| *ΔΔBcvic1/2-iii* | 839-5 *nit1 ΔBcvic1-ΔBcvic2::HPH* |
| Ectopic expression mutants (Complementation) |  |
| *ΔBcvic2-i + Bcvic2-1* | 839-5 NitM *ΔBcvic2::HPH Bcvic2+NAT* |
| *ΔΔBcvic1/2-i + Bcvic1-1/Bcvic2-1* | 839-5 NitM *ΔBcvic1-ΔBcvic2::HPH Bcvic1-1/Bcvic2-1+NAT* |
| *ΔΔBcvic1/2-i + Bcvic1-2/Bcvic2-2* | 839-5 NitM *ΔBcvic1-ΔBcvic2::HPH Bcvic1-2/Bcvic2-2+NAT* |
| *ΔΔBcvic1/2-i + Bcvic1-1* | 839-5 NitM *ΔBcvic1-ΔBcvic2::HPH Bcvic1-1+NAT* |
| *ΔΔBcvic1/2-i + Bcvic1-2* | 839-5 NitM *ΔBcvic1-ΔBcvic2::HPH Bcvic1-2+NAT* |
| *ΔΔBcvic1/2-i + Bcvic2-1* | 839-5 NitM *ΔBcvic1-ΔBcvic2::HPH Bcvic2-1+NAT* |
| Ectopic expression mutants (Hygromycin control) |  |
| 839-5 *nit1-HPH* | 839-5 *nit1 + HPH* |
| 839-5 NitM-*HPH* | 839-5 NitM + *HPH* |
| 839-1 *nit1-HPH* | 839-1 *nit1 + HPH* |
| 839-1 NitM-*HPH* | 839-1 NitM + *HPH* |

**S2 Table.** Similarity of candidate *vic* genes in the two VI bulks, vcg1 and vcg2.

| Candidate T4 gene models | Candidate B05-10 gene models | Putative function of the encoding protein | *vcg1* and *vcg2* nucleotide sequence identity (%) | *vcg1* and *vcg2* amino acid sequence identity (%) |
| --- | --- | --- | --- | --- |
| BofuT4_P145970<br>.1 | Bcin01g01190.<br>1 | putative aspartyl protease | 99.7 | 99.8 |
| BofuT4_P145980<br>.1 | Bcin01g01200 | No known domains | 99.3 | 99.5 |
| BofuT4_P145990<br>.1 | Bcin01g01210 | Syntaxin | 76.3* | 65.9* |
| BofuT4_P146010<br>.1 | Bcin01g01220 | Serine<br>esterase/NACHT/Ankyr<br>in repeats | 68.3* | 60.2* |
| BofuT4_P146020<br>.1 | Bcin01g01230 | tRNA methyl<br>transferase | 99.7 | 99.7 |
| BofuT4_P146070<br>.1 | Bcin01g01250 | No known domains | 97.4 | 97.9 |
| BofuT4_P146100<br>.1 | Bcin01p01290.<br>1 | No known domains | 99.4 | 99.8 |
| BofuT4_P146120<br>.1 | Bcin01p01300.<br>1 | oxidoreductase | 99.6 | 99.6 |
| BofuT4_P146130<br>.1 | Bcin01p01310.<br>1 | carboxypeptidase B | 98.4 | 98 |

\* Sequence comparisons only possible following PCR amplification and sequencing of genes from vcg1 since reads at the locus from the vcg1 bulk did not map to either the T4 or the B05.10 reference genomes.

**S3 Table.**
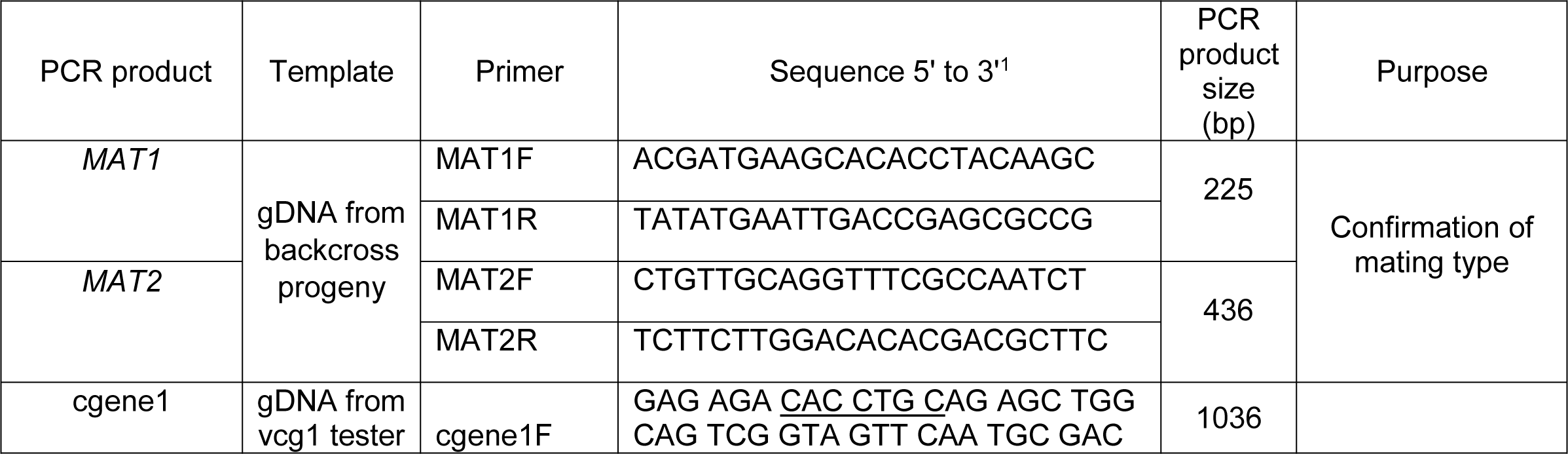
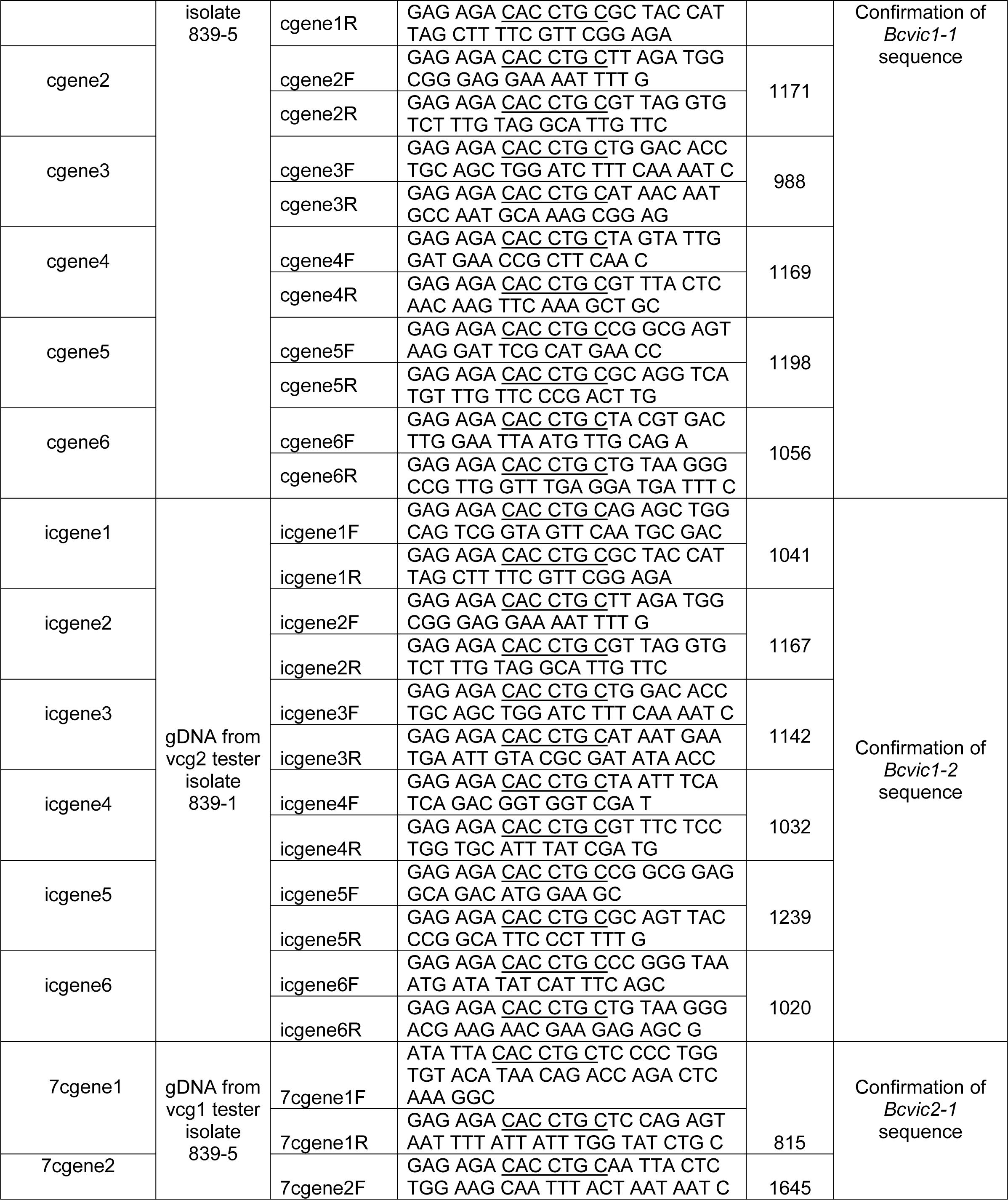
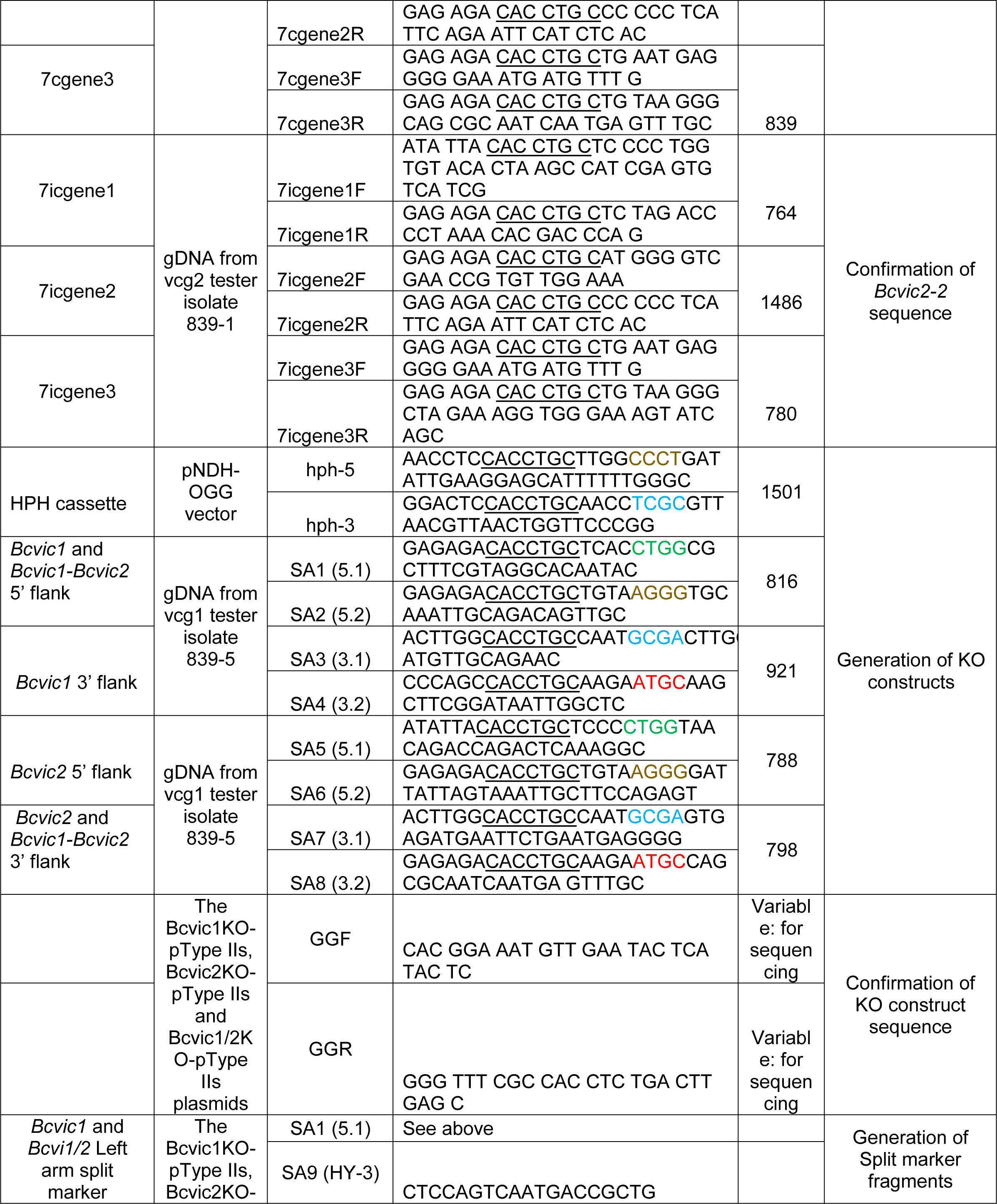
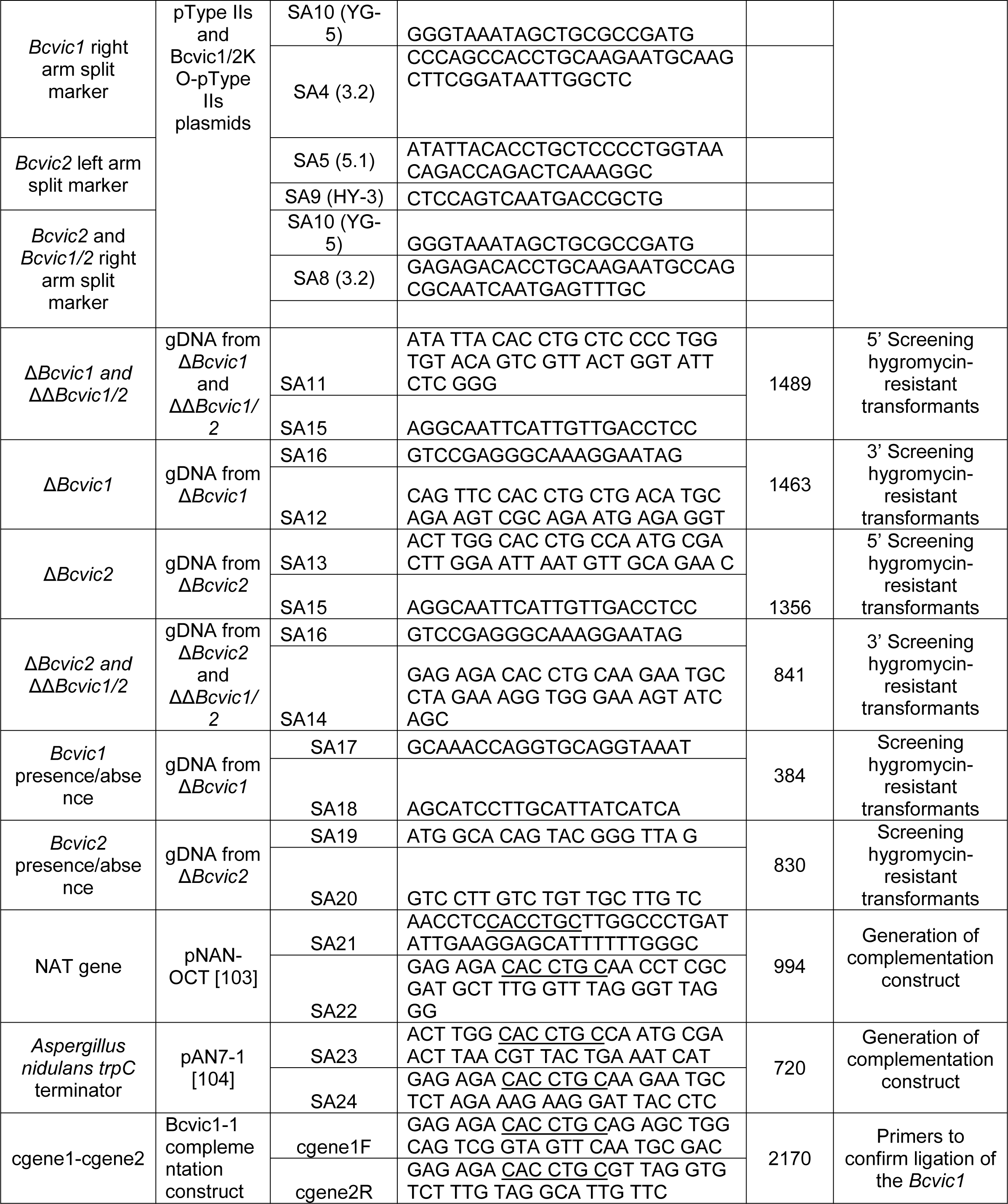
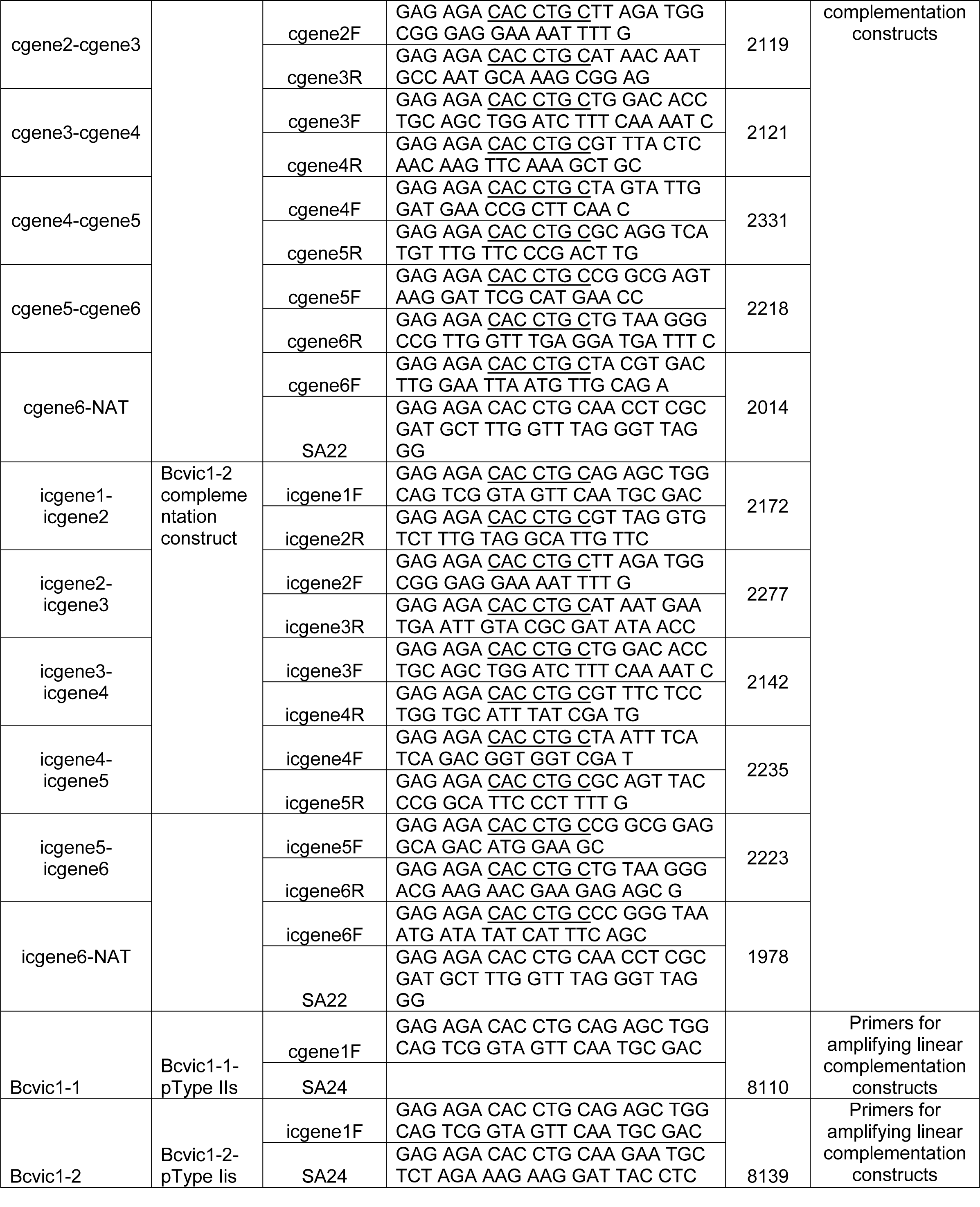
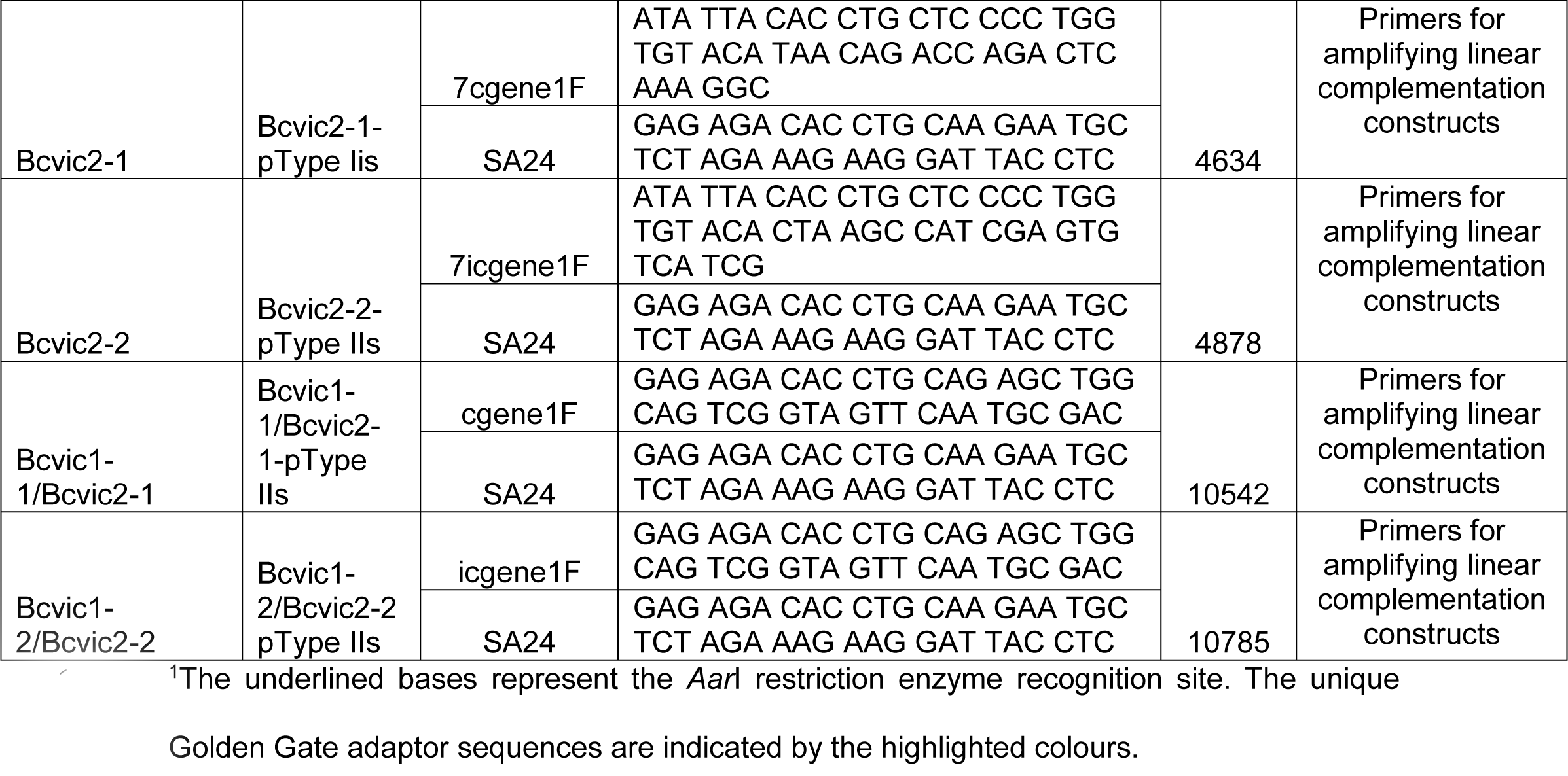
Primers used for PCR amplification.

## Supporting information figure captions

**S1 Fig.**
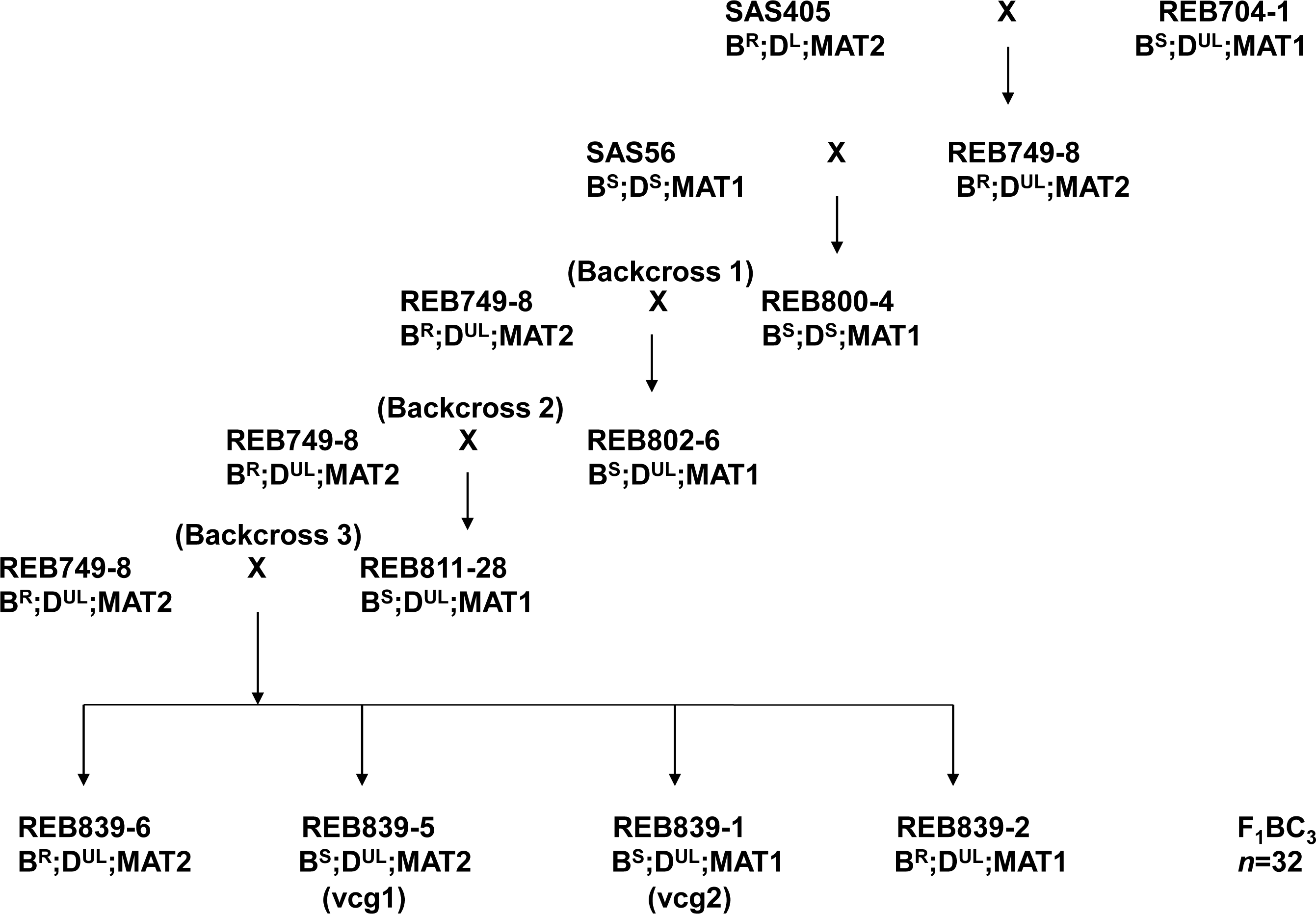
Pedigree of near-isogenic *Botrytis cinerea* strains. Thirty-two F1BC3 single ascospore isolates were generated from three backcrosses to the recurrent parent REB749-8. REB839-6, REB839-5, REB839-1 and REB839-2 are presented as four examples of progeny genotypes. REB839-5 and REB839-1 were used as the vcg1 (compatible with the recurrent parent REB749-8) and vcg2 (incompatible with the REB749-8) tester strains in this study. BR, benzimidazole-resistant; BS, benzimidazole-sensitive; DS, dicarboximide- sensitive; DL, low-level dicarboximide-resistant; DUL, ultra-low-level dicarboximide-resistant; MAT1, mating type 1; MAT2, mating type 2.

**S2 Fig.**
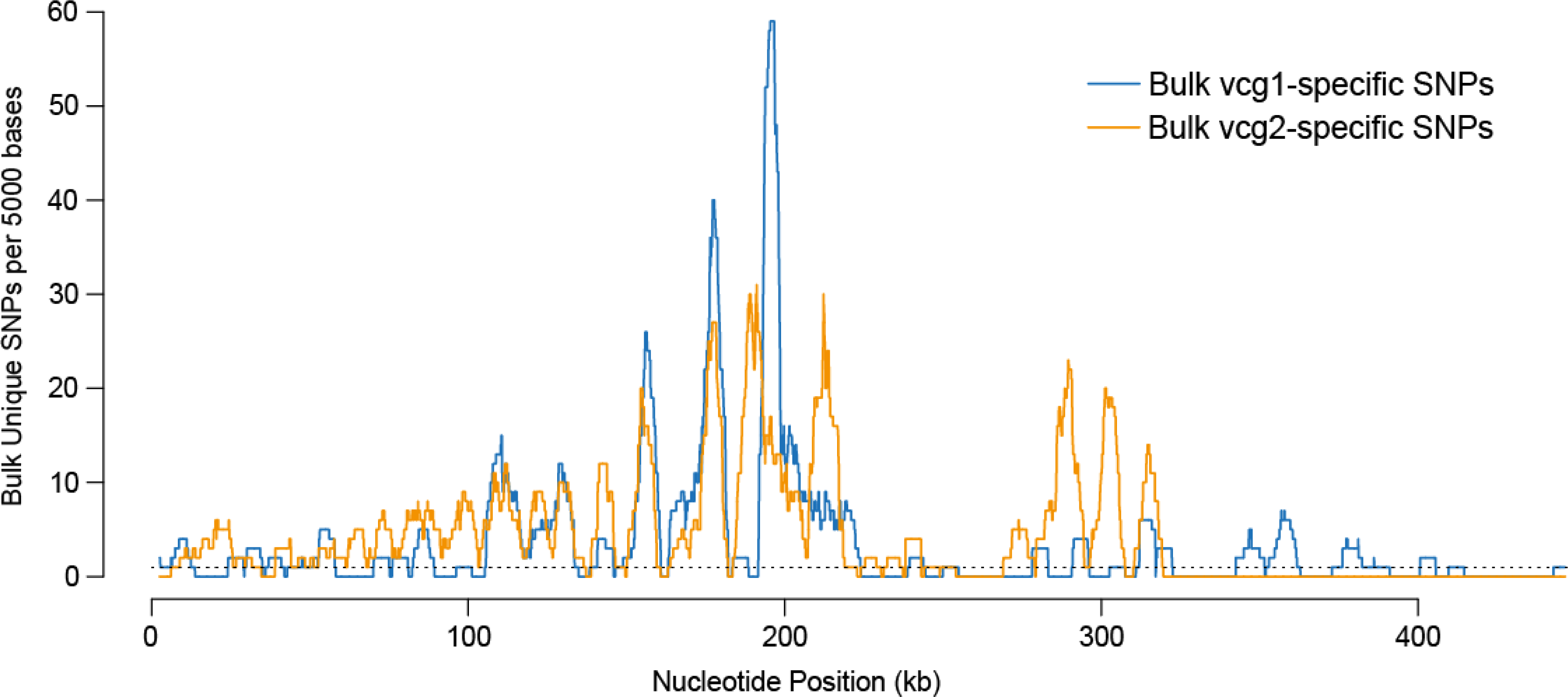
vcg1 and vcg2 bulk specific SNPs within scaffold bt4_SupSuperContig_110r_56_1 of the *Botrytis cinerea* T4 genome. A 5-kb sliding window with a 25-bp lag was applied to the entire scaffold, and the density of vcg1 and vcg2 bulk-specific SNPs is shown in blue and orange, respectively. The number of bulk-specific single nucleotide polymorphisms (SNPs) identified for this scaffold is substantially larger than for any other scaffold.

**S3 Fig.**
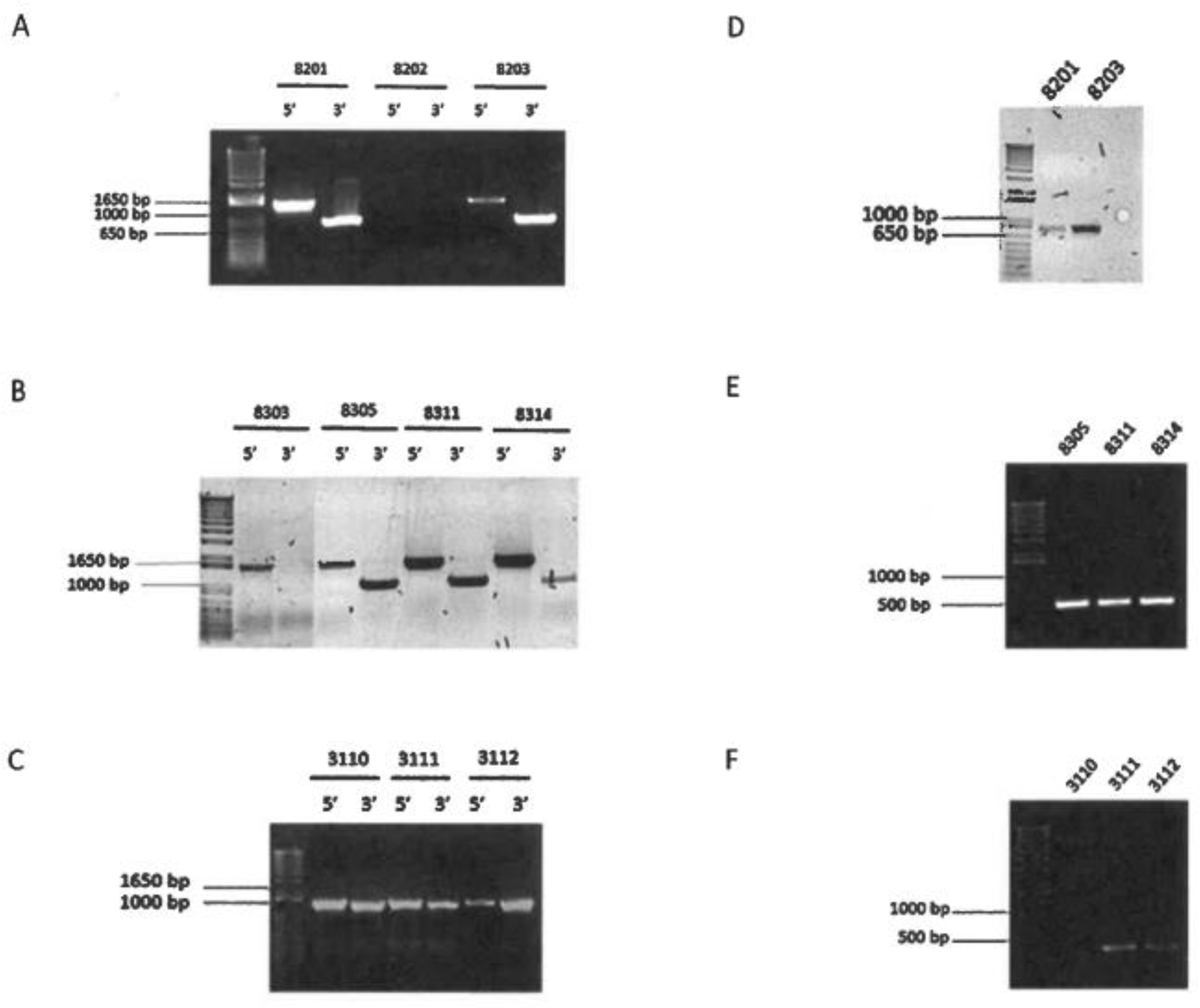
Polymerase Chain Reaction (PCR) screen of *Botrytis cinerea* gene knockout mutants. (A-C) Homologous recombination at the 5’ and 3’ flanks in a subset of independent *Bcvic2*, *Bcvic1/Bcvic2* and *Bcvic1* recombinants, respectively. (D-F) Detection of the native *Bcvic2* and *Bcvic1* genes in *Botrytis cinerea Bcvic2*, *Bcvic1/Bcvic2* and *Bcvic1* recombinants, respectively.

**S4 Fig.**
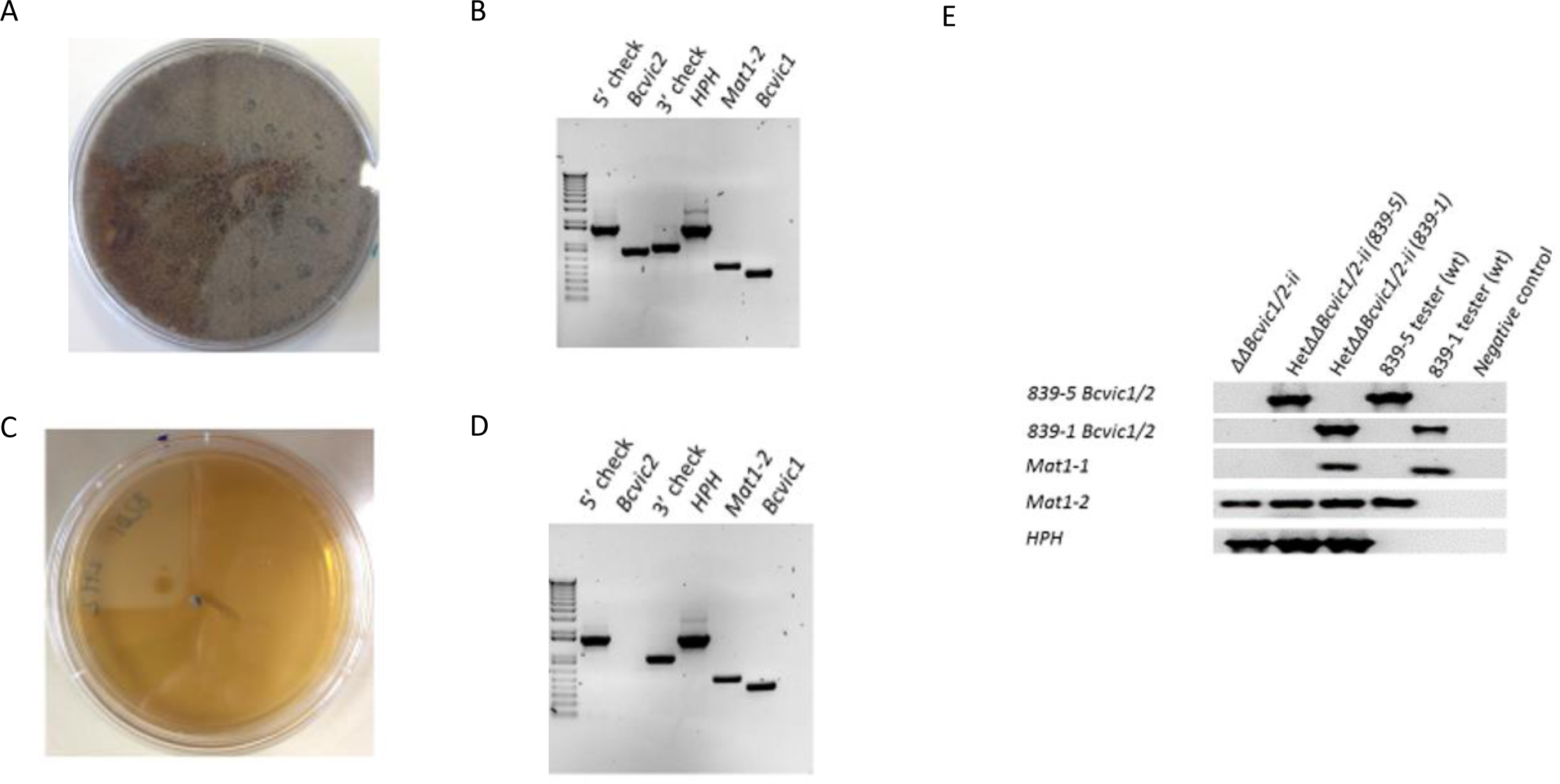
Growth morphology and Polymerase Chain Reaction (PCR)analysis of fast and slow growing HetΔBcvic2 single-spore isolates. (A, B) Fast-growing isolate. (C, D) Slow- growing isolate. (E) PCR analysis of slow-growing *HetΔΔBcvic1/2* single spore isolate. WT, wild-type; *HPH*, hygromycin B phosphotransferase.

**S5 Fig.**
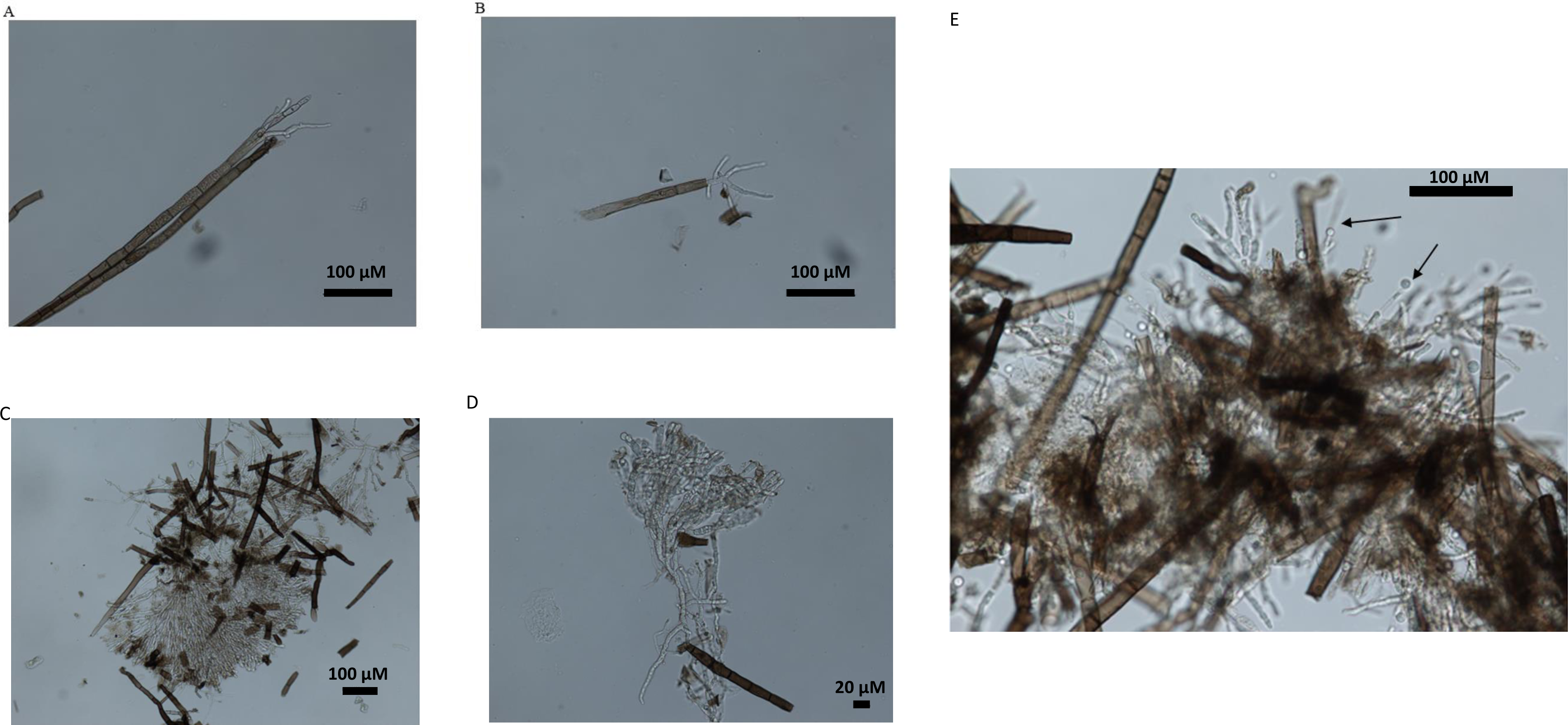
Hyphal regeneration and protoplasting of melanised *ΔBcvic2* and *ΔΔBcvic1/2* mutants. (A, B) Hyphal regeneration 3 d post-inoculation. (A) *ΔΔBcvic1/2* and (B) *ΔBcvic2*. (C, D) Hyphal regeneration 5 days post-inoculation. (C) *ΔΔBcvic1/2* and (D) *ΔBcvic2*. (E) Protoplasts generated from macerated mycelia. Black arrows point towards budding protoplasts. Scale bar is 100 µM.

**S6 Fig.**
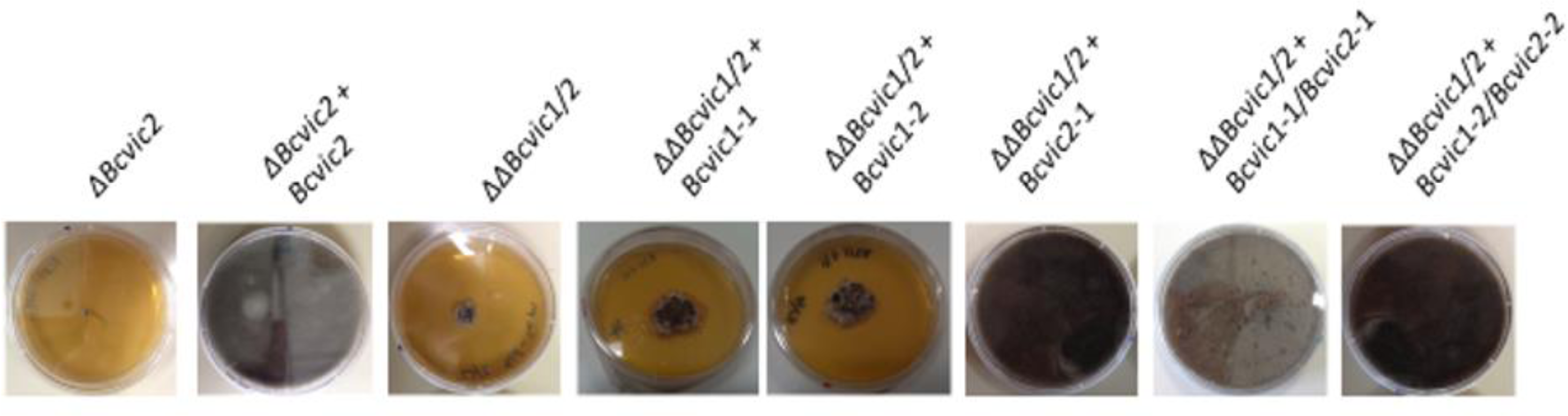
Growth phenotype of *ΔBcvic2* and *ΔΔBcvic1/2* knockout and complementation mutants. Single spore isolates of the various mutants were inoculated on the centre of an agar plate. Photographs were taken 8 days post-inoculation. (A-H) *ΔBcvic2*, *ΔBcvic2+Bcvic2, ΔΔBcvic1/2*, *ΔΔBcvic1/2+Bcvic1-1*, *ΔΔBcvic1/2+Bcvic2-1*, *ΔΔBcvic1/2+Bcvic1-1/Bcviv2-1 and ΔΔBcvic1/2+Bcvic1-2/Bcvic2-2*, respectively.

## References

1. Boswell GP, Hopkins S. Linking hyphal growth to colony dynamics: spatially explicit models of mycelia. Fungal Ecology. 2008;1:143–54.

2. Richard F, Glass NL, Pringle A. Cooperation among germinating spores facilitates the growth of the fungus, *Neurospora crassa*. Biology Letters. 2012;8:419–22.

3. Bastiaans E, Debets AJM, Aanen DK. Experimental demonstration of the benefits of somatic fusion and the consequences for allorecognition. Evolution. 2015;69:1091–9.

4. Fischer MS, Glass NL. Communicate and fuse: how filamentous fungi establish and maintain an interconnected mycelial network. Frontiers in Microbiology. 2019;10:619.

5. Bastiaans E, Aanen DK, Debets AJ, Hoekstra RF, Lestrade B, Maas MF. Regular bottlenecks and restrictions to somatic fusion prevent the accumulation of mitochondrial defects in *Neurospora*. Philosophical Transactions of the Royal Society B. 2014;369:20130448.

6. Debets F, Yang X, Griffiths AJ. Vegetative incompatibility in *Neurospora*: its effect on horizontal transfer of mitochondrial plasmids and senescence in natural populations. Current Genetics. 1994;26:113–9.

7. Nuss DL. Hypovirulence: mycoviruses at the fungal-plant interface. Nature Reviews Microbiology. 2005;3:632–42.

8. Son M, Yu J, Kim K-H. Five questions about mycoviruses. PLoS Pathogens. 2015;11:e1005172.

9. Ghabrial SA, Castón JR, Jiang D, Nibert ML, Suzuki N. 50-plus years of fungal viruses. Virology. 2015;479–480:356-68.

10. Gonçalves AP, Glass NL. Fungal social barriers: to fuse, or not to fuse, that is the question. Communicative & Integrative Biology. 2020;13:39–42.

11. Heller J, Zhao J, Rosenfield G, Kowbel DJ, Gladieux P, Glass NL. Characterization of greenbeard genes involved in long-distance kind discrimination in a microbial eukaryote. PLoS Biology. 2016;14:e1002431.

12. Gonçalves AP, Heller J, Span EA, Rosenfield G, Do HP, Palma-Guerrero J, et al. Allorecognition upon fungal cell-cell contact determines social cooperation and impacts the acquisition of multicellularity. Current Biology. 2019;29:3006–17.e3.

13. Heller J, Clavé C, Gladieux P, Saupe SJ, Glass NL. NLR surveillance of essential SEC-9 SNARE proteins induces programmed cell death upon allorecognition in filamentous fungi. Proceedings of the National Academy of Sciences U S A. 2018;115:E2292–E301.

14. Daskalov A, Gladieux P, Heller J, Glass NL. Programmed cell death in *Neurospora crassa* is controlled by the allorecognition determinant *rcd-1*. Genetics. 2019;213:1387–400.

15. Dyer PS. Self/non-self recognition: microbes playing hard to get. Current Biology. 2019;29:R866–R8.

16. Paoletti M. Vegetative incompatibility in fungi: From recognition to cell death, whatever does the trick. Fungal Biology Reviews. 2016;30:152–62.

17. Glass NL, Dementhon K. Non-self recognition and programmed cell death in filamentous fungi. Current Opinion in Microbiology. 2006;9:553–8.

18. Gonçalves AP, Heller J, Daskalov A, Videira A, Glass NL. Regulated forms of cell death in fungi. Frontiers in Microbiology. 2017;8:1837.

19. Garnjobst L, Wilson JF. Heterocaryosis and protoplasmic incompatibility in *Neurospora crassa*. Proceedings of the National Academy of Sciences U S A. 1956;42:613–8.

20. Saupe SJ. Molecular genetics of heterokaryon incompatibility in filamentous ascomycetes. Microbiology and Molecular Biology Reviews. 2000;64:489–502.

21. Leslie JF. Fungal vegetative compatibility. Annual Review of Phytopathology. 1993;31:127–50.

22. Muirhead CA, Glass NL, Slatkin M. Multilocus self-recognition systems in fungi as a cause of trans-species polymorphism. Genetics. 2002;161:633–41.

23. Glass NL, Kaneko I. Fatal attraction: nonself recognition and heterokaryon incompatibility in filamentous fungi. Eukaryotic Cell. 2003;2:1–8.

24. Pál K, van Diepeningen AD, Varga J, Hoekstra RF, Dyer PS, Debets AJM. Sexual and vegetative compatibility genes in the aspergilli. Studies in Mycology. 2007;59:19–30.

25. Zhao J, Gladieux P, Hutchison E, Bueche J, Hall C, Perraudeau F, et al. Identification of allorecognition loci in *Neurospora crassa* by genomics and evolutionary approaches. Molecular Biology and Evolution. 2015;32:2417–32.

26. Worrall JJ. Somatic incompatibility in Basidiomycetes. Mycologia. 1997;89:24–36.

27. Giovannetti M, Sbrana C, Strani P, Agnolucci M, Rinaudo V, Avio L. Genetic diversity of isolates of *Glomus mosseae* from different geographic areas detected by vegetative compatibility testing and biochemical and molecular analysis. Applied and Environmental Microbiology. 2003;69:616–24.

28. Daskalov A, Heller J, Herzog S, Fleißner A, Glass NL. Molecular mechanisms regulating cell fusion and heterokaryon formation in filamentous fungi. Microbiology Spectrum. 2017;5:10.1128/microbiolspec.FUNK-0015-2016

29. Kaneko I, Dementhon K, Xiang Q, Glass NL. Nonallelic interactions between *het-c* and a polymorphic locus, *pin-c*, are essential for nonself recognition and programmed cell death in *Neurospora crassa*. Genetics. 2006;172:1545–55.

30. Espagne E, Balhadère P, Penin ML, Barreau C, Turcq B. HET-E and HET-D belong to a new subfamily of WD40 proteins involved in vegetative incompatibility specificity in the fungus *Podospora anserina*. Genetics. 2002;161:71–81.

31. Saupe S, Descamps C, Turcq B, Begueret J. Inactivation of the *Podospora anserina* vegetative incompatibility locus *het-c*, whose product resembles a glycolipid transfer protein, drastically impairs ascospore production. Proceedings of the National Academy of Sciences U S A. 1994;91:5927–31.

32. Saupe S, Turcq B, Bégueret J. A gene responsible for vegetative incompatibility in the fungus *Podospora anserina* encodes a protein with a GTP-binding motif and Gβ homologous domain. Gene. 1995;162:135–9.

33. Paoletti M, Clavé C. The fungus-specific HET domain mediates programmed cell death in *Podospora anserina*. Eukaryotic Cell. 2007;6:2001–8.

34. Daskalov A, Habenstein B, Martinez D, Debets AJM, Sabaté R, Loquet A, et al. Signal transduction by a fungal NOD-like receptor based on propagation of a prion amyloid fold. PLoS Biology. 2015;13:e1002059.

35. Turcq B, Deleu C, Denayrolles N, Bégueret J. Two allelic genes responsible for vegetative incompatibility in the fungus *Podospora anserina* are not essential for cell viabilty. Molecular and General Genetics. 1991;228:265–9.

36. Balguerie A, Dos Reis S, Ritter C, Chaignepain S, Coulary-Salin B, Forge V, et al. Domain organization and structure-function relationship of the HET-s prion protein of *Podospora anserina*. The EMBO Journal. 2003;22:2071–81.

37. Greenwald J, Buhtz C, Ritter C, Kwiatkowski W, Choe S, Maddelein M-L, et al. The mechanism of prion inhibition by HET-S. Molecular Cell. 2010;38:889–99.

38. Mathur V, Seuring C, Riek R, Saupe SJ, Liebman SW. Localization of HET-S to the cell periphery, not to [Het-s] aggregates, is associated with [Het-s]-HET-S toxicity. Molecular and Cell Biology. 2012;32:139–53.

39. Seuring C, Greenwald J, Wasmer C, Wepf R, Saupe SJ, Meier BH, et al. The mechanism of toxicity in HET-S/HET-s prion incompatibility. PLoS Biology. 2012;10:e1001451.

40. Bidard F, Clavé C, Saupe SJ. The transcriptional response to nonself in the fungus *Podospora anserina*. G3 Genes Genomes Genetics. 2013;3:1015–30.

41. Lamacchia M, Dyrka W, Breton A, Saupe SJ, Paoletti M. Overlapping *Podospora anserina* transcriptional responses to bacterial and fungal non self indicate a multilayered innate immune response. Frontiers in Microbiology. 2016;7:471-.

42. Pinan-Lucarré B, Paoletti M, Clavé C. Cell death by incompatibility in the fungus *Podospora*. Seminars in Cancer Biology. 2007;17:101–11.

43. Belov AA, Witte TE, Overy DP, Smith ML. Transcriptome analysis implicates secondary metabolite production, redox reactions, and programmed cell death during allorecognition in *Cryphonectria parasitica*. G3 Genes Genomes Genetics. 2021;11.

44. Witte TE, Shields S, Heberlig GW, Darnowski MG, Belov A, Sproule A, et al. A metabolomic study of vegetative incompatibility in *Cryphonectria parasitica*. Fungal Genetics and Biology. 2021;157:103633.

45. García-Pedrajas MD, Cañizares MC, Sarmiento-Villamil JL, Jacquat AG, Dambolena JS. Mycoviruses in biological control: from basic research to field implementation. Phytopathology. 2019;109:1828–39.

46. Williamson B, Tudzynski B, Tudzynski P, van Kan JA. *Botrytis cinerea*: the cause of grey mould disease. Molecular Plant Pathology. 2007;8:561–80.

47. Staats M, Kan JALv. Genome update of *Botrytis cinerea* strains B05.10 and T4. Eukaryotic Cell. 2012;11:1413–4.

48. Choquer M, Rascle C, Gonçalves IR, de Vallée A, Ribot C, Loisel E, et al. The infection cushion of *Botrytis cinerea*: a fungal ‘weapon’ of plant-biomass destruction. Environmental Microbiology. 2021;23:2293–314.

49. Hao F, Ding T, Wu M, Zhang J, Yang L, Chen W, et al. Two novel hypovirulence-associated mycoviruses in the phytopathogenic fungus *Botrytis cinerea*: molecular characterization and suppression of infection cushion formation. Viruses. 2018;10:254.

50. Castro M, Kramer K, Valdivia L, Ortiz S, Castillo A. A double-stranded RNA mycovirus confers hypovirulence-associated traits to *Botrytis cinerea*. FEMS Microbiology Letters. 2003;228:87–91.

51. Potgieter CA, Castillo A, Castro M, Cottet L, Morales A. A wild-type *Botrytis cinerea* strain co- infected by double-stranded RNA mycoviruses presents hypovirulence-associated traits. Virology Journal. 2013;10:220.

52. Zhang D-X, Nuss DL. Engineering super mycovirus donor strains of chestnut blight fungus by systematic disruption of multilocus *vic* genes. Proceedings of the National Academy of Sciences U S A. 2016;113:2062–7.

53. Stauder CM, Nuss DL, Zhang D-X, Double ML, MacDonald WL, Metheny AM, et al. Enhanced hypovirus transmission by engineered super donor strains of the chestnut blight fungus, *Cryphonectria parasitica*, into a natural population of strains exhibiting diverse vegetative compatibility genotypes. Virology. 2019;528:1–6.

54. Beever RE, Weeds PL. Taxonomy and genetic variation of Botrytis and Botryotinia. In: Elad Y, Williamson B, Tudzynski P, Delen N, editors. Botrytis: biology, pathology and control. Dordrecht, The Netherlands: Springer; 2007. p. 29-52.

55. Liu J-J, Sniezko RA, Zamany A, Williams H, Omendja K, Kegley A, et al. Comparative transcriptomics and RNA-seq-based bulked segregant analysis reveals genomic basis underlying *Cronartium ribicola vcr2* virulence. Frontiers in Microbiology. 2021;12.

56. Plissonneau C, Benevenuto J, Mohd-Assaad N, Fouché S, Hartmann FE, Croll D. Using population and comparative genomics to understand the genetic basis of effector-driven fungal pathogen evolution. Frontiers in Plant Science. 2017;8:119.

57. Wu JQ, Sakthikumar S, Dong C, Zhang P, Cuomo CA, Park RF. Comparative genomics integrated with association analysis identifies candidate effector genes corresponding to *Lr20* in phenotype-paired *Puccinia triticina* isolates from Australia. Frontiers in Plant Science. 2017;8:148.

58. Acosta Morel W, Anta Fernández F, Baroncelli R, Becerra S, Thon MR, van Kan JAL, et al. A major effect gene controlling development and pathogenicity in *Botrytis cinerea* identified through genetic analysis of natural mycelial non-pathogenic isolates. Frontiers in Plant Science. 2021;12:663870.

59. Beever R, Parkes S. Use of nitrate non-utilising (Nit) mutants to determine vegetative compatibility in *Botryotinia fuckeliana* (*Botrytis cinerea*). European Journal of Plant Pathology. 2003;109:607–13.

60. Choi GH, Dawe AL, Churbanov A, Smith ML, Milgroom MG, Nuss DL. Molecular characterization of vegetative incompatibility genes that restrict hypovirus transmission in the chestnut blight fungus *Cryphonectria parasitica*. Genetics. 2012;190:113–27.

61. Chevanne D, Bastiaans E, Debets A, Saupe SJ, Clavé C, Paoletti M. Identification of the *het-r* vegetative incompatibility gene of *Podospora anserina* as a member of the fast evolving *HNWD* gene family. Current Genetics. 2009;55:93–102.

62. Li J, Mahajan A, Tsai MD. Ankyrin repeat: a unique motif mediating protein-protein interactions. Biochemistry. 2006;45:15168–78.

63. Zeytuni N, Zarivach R. Structural and functional discussion of the tetra-trico-peptide repeat, a protein interaction module. Structure. 2012;20:397–405.

64. Daskalov A, Mitchell PS, Sandstrom A, Vance RE, Glass NL. Molecular characterization of a fungal gasdermin-like protein. Proceedings of the National Academy of Sciences U S A. 2020;117:18600–7.

65. Roudaire T, Héloir M-C, Wendehenne D, Zadoroznyj A, Dubrez L, Poinssot B. Cross kingdom immunity: the role of immune receptors and downstream signaling in animal and plant cell death. Frontiers in Immunology. 2021;11:612452.

66. Burdett H, Bentham AR, Williams SJ, Dodds PN, Anderson PA, Banfield MJ, et al. The plant “resistosome”: structural insights into immune signaling. Cell Host & Microbe. 2019;26:193–201.

67. Zheng D, Liwinski T, Elinav E. Inflammasome activation and regulation: toward a better understanding of complex mechanisms. Cell Discovery. 2020;6:36.

68. Wang J, Hu M, Wang J, Qi J, Han Z, Wang G, et al. Reconstitution and structure of a plant NLR resistosome conferring immunity. Science. 2019;364:eaav5870.

69. Wang J, Wang J, Hu M, Wu S, Qi J, Wang G, et al. Ligand-triggered allosteric ADP release primes a plant NLR complex. Science. 2019;364:eaav5868.

70. Bi G, Su M, Li N, Liang Y, Dang S, Xu J, et al. The ZAR1 resistosome is a calcium-permeable channel triggering plant immune signaling. Cell. 2021;184:3528–41.e12.

71. Han J, Pluhackova K, Böckmann RA. The multifacted role of SNARE proteins in membrane fusion. Frontiers in Physiology. 2017;8:5.

72. Pratelli R, Sutter J-U, Blatt MR. A new catch in the SNARE. Trends in Plant Science. 2004;9:187–95.

73. Kloepper TH, Kienle CN, Fasshauer D. An elaborate classification of SNARE proteins sheds light on the conservation of the eukaryotic endomembrane system. Molecular Biology of the Cell. 2007;18:3463–71.

74. Qi Z, Liu M, Dong Y, Zhu Q, Li L, Li B, et al. The syntaxin protein (MoSyn8) mediates intracellular trafficking to regulate conidiogenesis and pathogenicity of rice blast fungus. New Phytologist. 2016;209:1655–67.

75. Song W, Dou X, Qi Z, Wang Q, Zhang X, Zhang H, et al. R-SNARE homolog MoSec22 is required for conidiogenesis, cell wall integrity, and pathogenesis of *Magnaporthe oryzae*. PLoS ONE. 2010;5:e13193.

76. Dou X, Wang Q, Qi Z, Song W, Wang W, Guo M, et al. MoVam7, a conserved SNARE involved in vacuole assembly, is required for growth, endocytosis, ROS accumulation, and pathogenesis of *Magnaporthe oryzae*. PLoS ONE. 2011;6:e16439.

77. Kienle N, Kloepper TH, Fasshauer D. Phylogeny of the SNARE vesicle fusion machinery yields insights into the conservation of the secretory pathway in fungi. BMC Evolutionary Biology. 2009;9:19.

78. Sutton RB, Fasshauer D, Jahn R, Brunger AT. Crystal structure of a SNARE complex involved in synaptic exocytosis at 2.4 Å resolution. Nature. 1998;395:347–53.

79. Koike S, Jahn R. SNARE proteins: zip codes in vesicle targeting? Biochemistry Journal. 2022;479:273–88.

80. Ko Y-H, So K-K, Kim J-M, Kim D-H. Heterokaryon analysis of a Cdc48-like gene, CpCdc48, from the chestnut blight fungus Cryphonectria parasitica demonstrates it is essential for cell division and growth. Fungal Genetics and Biology. 2016;88:1–12.

81. Wang M, Jin H. Spray-induced gene silencing: a powerful innovative strategy for crop protection. Trends in Microbiology. 2017;25:4–6.

82. Wang M, Weiberg A, Lin F-M, Thomma BPHJ, Huang H-D, Jin H. Bidirectional cross-kingdom RNAi and fungal uptake of external RNAs confer plant protection. Nat Plants. 2016;2:16151.

83. Khalifa ME, MacDiarmid RM. A mechanically transmitted DNA mycovirus Is targeted by the defence machinery of its host, *Botrytis cinerea*. Viruses. 2021;13:1315.

84. Amselem J, Cuomo CA, van Kan JA, Viaud M, Benito EP, Couloux A, et al. Genomic analysis of the necrotrophic fungal pathogens *Sclerotinia sclerotiorum* and *Botrytis cinerea*. PLoS Genetics. 2011;7:e1002230.

85. Vogel HJ. A convenient growth medium for *Neurospora crassa*. Microbial Genetics Bulletin. 1956;13:42–7.

86. Möller EM, Bahnweg G, Sandermann H, Geiger HH. A simple and efficient protocol for isolation of high molecular weight DNA from filamentous fungi, fruit bodies, and infected plant tissues. Nucleic Acids Research. 1992;20:6115–6.

87. Faretra F, Antonacci E. Production of apothecia of *Botryotinia fuckeliana* (de Bary) Whetz. under controlled environmental conditions. Phytopathologia Mediterranea. 1987;26:29–35.

88. Faretra F, Antonacci E, Pollastro S. Sexual behaviour and mating system of *Botryotinia fuckeliana*, teleomorph of *Botrytis cinerea*. Journal of General Microbiology. 1988;134:2543–50.

89. Cox MP, Peterson DA, Biggs PJ. SolexaQA: At-a-glance quality assessment of Illumina second- generation sequencing data. BMC Bioinformatics. 2010;11:485-.

90. Li H, Durbin R. Fast and accurate long-read alignment with Burrows-Wheeler transform. Bioinformatics. 2010;26:589–95.

91. Robinson JT, Thorvaldsdóttir H, Winckler W, Guttman M, Lander ES, Getz G, et al. Integrative genomics viewer. Nature Biotechnology. 2011;29:24–6.

92. Li H, Handsaker B, Wysoker A, Fennell T, Ruan J, Homer N, et al. The Sequence Alignment/Map format and SAMtools. Bioinformatics. 2009;25:2078–9.

93. Danecek P, Auton A, Abecasis G, Albers CA, Banks E, DePristo MA, et al. The variant call format and VCFtools. Bioinformatics. 2011;27:2156–8.

94. Foissac S, Gouzy J, Rombauts S, Mathe C, Amselem J, Sterck L, et al. Genome annotation in plants and fungi: EuGene as a model platform. Bioinformatics. 2008;3:87–97.

95. Salamov AA, Solovyev VV. Ab initio gene finding in *Drosophila* genomic DNA. Genome Research. 2000;10:516–22.

96. Solovyev V, Kosarev P, Seledsov I, Vorobyev D. Automatic annotation of eukaryotic genes, pseudogenes and promoters. Genome Biology. 2006;7:S10.

97. Van Kan JA, Stassen JH, Mosbach A, Van Der Lee TA, Faino L, Farmer AD, et al. A gapless genome sequence of the fungus *Botrytis cinerea*. Molecular Plant Pathology. 2017;18:75–89.

98. Mistry J, Chuguransky S, Williams L, Qureshi M, Salazar GA, Sonnhammer ELL, et al. Pfam: The protein families database in 2021. Nucleic Acids Research. 2021;49:D412–D9.

99. Finn RD, Coggill P, Eberhardt RY, Eddy SR, Mistry J, Mitchell AL, et al. The Pfam protein families database: towards a more sustainable future. Nucleic Acids Research. 2016;44:D279–85.

100. Madeira F, Park YM, Lee J, Buso N, Gur T, Madhusoodanan N, et al. The EMBL-EBI search and sequence analysis tools APIs in 2019. Nucleic Acids Research. 2019;47:W636–w41.

101. Engler C, Gruetzner R, Kandzia R, Marillonnet S. Golden Gate shuffling: A one-pot DNA shuffling method based on Type IIs restriction enzymes. PLoS ONE. 2009;4:e5553.

102. Engler C, Kandzia R, Marillonnet S. A one pot, one step, precision cloning method with high throughput capability. PLoS One. 2008;3:e3647.

103. Schumacher J. Tools for *Botrytis cinerea*: New expression vectors make the gray mold fungus more accessible to cell biology approaches. Fungal Genetics and Biology. 2012;49:483–97.

104. Punt PJ, Oliver RP, Dingemanse MA, Pouwels PH, van den Hondel CA. Transformation of *Aspergillus* based on the hygromycin B resistance marker from *Escherichia coli*. Gene. 1987;56:117–24.

105. ten Have A, Mulder W, Visser J, van Kan JA. The endopolygalacturonase gene *Bcpg1* is required for full virulence of *Botrytis cinerea*. Molecular Plant-Microbe Interactions. 1998;11:1009–16.

